# Spatially resolved cell atlas of the teleost telencephalon and deep homology of the vertebrate forebrain

**DOI:** 10.1101/2023.07.20.549873

**Authors:** Brianna E. Hegarty, George W. Gruenhagen, Zachary V. Johnson, Cristina M. Baker, Jeffrey T. Streelman

## Abstract

The telencephalon has undergone remarkable diversification and expansion throughout vertebrate evolution, exhibiting striking differences in structural and functional complexity. Nevertheless, fundamental features are shared across vertebrate taxa, such as the presence of distinct regions including the pallium, subpallium, and olfactory structures. Teleost fishes have a uniquely ‘everted’ telencephalon, which has made it challenging to compare brain regions in fish to those in other vertebrates. Here we combine spatial transcriptomics and single-nucleus RNA-sequencing to generate a spatially-resolved transcriptional atlas of the cichlid fish telencephalon. We then compare cell-types and anatomical regions in the cichlid telencephalon with those in amphibians, reptiles, birds, and mammals. We uncover striking transcriptional similarities between cell populations in the fish telencephalon and subpallial, hippocampal, and cortical cell populations in tetrapods. Ultimately, our work lends new insights into the organization and evolution of conserved cell-types and regions in the vertebrate forebrain.

## INTRODUCTION

The forebrain houses regions involved in advanced cognition such as social behavior, memory and learning across vertebrate lineages. Despite its conserved functions, this structure has grown larger and increasingly complex over evolutionary time, exhibiting marked variations across vertebrate taxa including the mammalian six-layered neocortex, the DVR in sauropsids (reptiles and birds), and the ‘everted’ teleost pallium of ray-finned fish (Actinopterygii). As a result, the evolutionary relationships of specific forebrain regions across species are still actively debated.

Several subregions of the teleost telencephalon have been compared to mammalian brain regions involved in regulating social behaviors, including the hippocampus, striatum and septum^1^. However, establishing homologues between telencephalic subdivisions in fish to those of distantly-related vertebrate clades has been historically challenging, due in part to the ‘everted’ morphology of the teleost telencephalon. The unique outward folding of the teleost pallium during development, known as eversion, leads to an altered arrangement of pallial zones compared to the ‘evaginated’ brains of other vertebrate lineages^2, 3^. Multiple proposed models of eversion attempt to identify the counterparts of other vertebrate pallial territories in fish^2–6^. However, these models are unresolved, leaving the evolutionary identities of most divisions of the teleost pallium in question. Modern omics technologies capable of simultaneously profiling thousands of genes are powerful tools to address these long-standing questions. Recent studies in different tetrapod lineages have begun to utilize these techniques to investigate vertebrate brain evolution at an unprecedented resolution^7–12^. Advances in methods for comparative analysis of these data have further increased the ability to establish homologous cell-types across distant phyla^13, 14^.

We recently employed single-nucleus RNA-sequencing (snRNA-seq) to link genomic signatures of behavioral evolution to specific cell populations in the cichlid telencephalon^15^. Here, we perform spatial transcriptomics (ST) and map cell populations sequenced at single-cell resolution to create a spatially resolved transcriptional atlas of the cichlid telencephalon. We then survey the cellular architecture of this critical brain structure, and utilize comparative approaches spanning all five major vertebrate lineages to generate new and unanticipated insights into the evolution of cell-types and neuroanatomical regions in the teleost forebrain.

## RESULTS

### Spatially resolved gene expression profiles in the the teleost brain

To investigate the anatomical landscape of the adult *Mchenga conophoros* (MC) telencephalon, we generated ST data using the 10X Genomics Visium platform (Fig. 1A). We dissected the telencephala from two adult male MCs (Table S1), collecting 3-4 representative 10μm coronal sections along the rostrocaudal axis per subject on the individual capture areas of 10X Visium slides (Fig. 1B, Fig. S1). Due to rostrocaudal variation between the left and right telencephalic hemispheres on individual capture areas, we visualized and analyzed each hemisphere separately (Fig. 1B). In total, >500 million RNA reads (68±2 million reads per capture area) were sequenced and aligned to the cichlid *Maylandia zebra* reference genome^16^. A total of 3,971 spots (567±65 spots per capture area) contained tissue and at least 1 unique molecular identifier (UMI), averaging 9,729±81 UMIs (range: 73-37,124) and 3,639±20 genes per spot (range: 73-7,263; Fig. S2). Clustering parameters were chosen systematically using ChooseR, which evaluates clustering quality based on robustness metrics of bootstraps (Fig. S3; Table S2). Using these near optimal parameters, 33 unbiased clusters of spots were formed (Fig. 1C-D). Marker genes of unbiased clusters were determined (Table S3) and among these were canonical markers of well-defined anatomical regions in the teleost telencephalon (Fig. S4).

**Figure 1.**
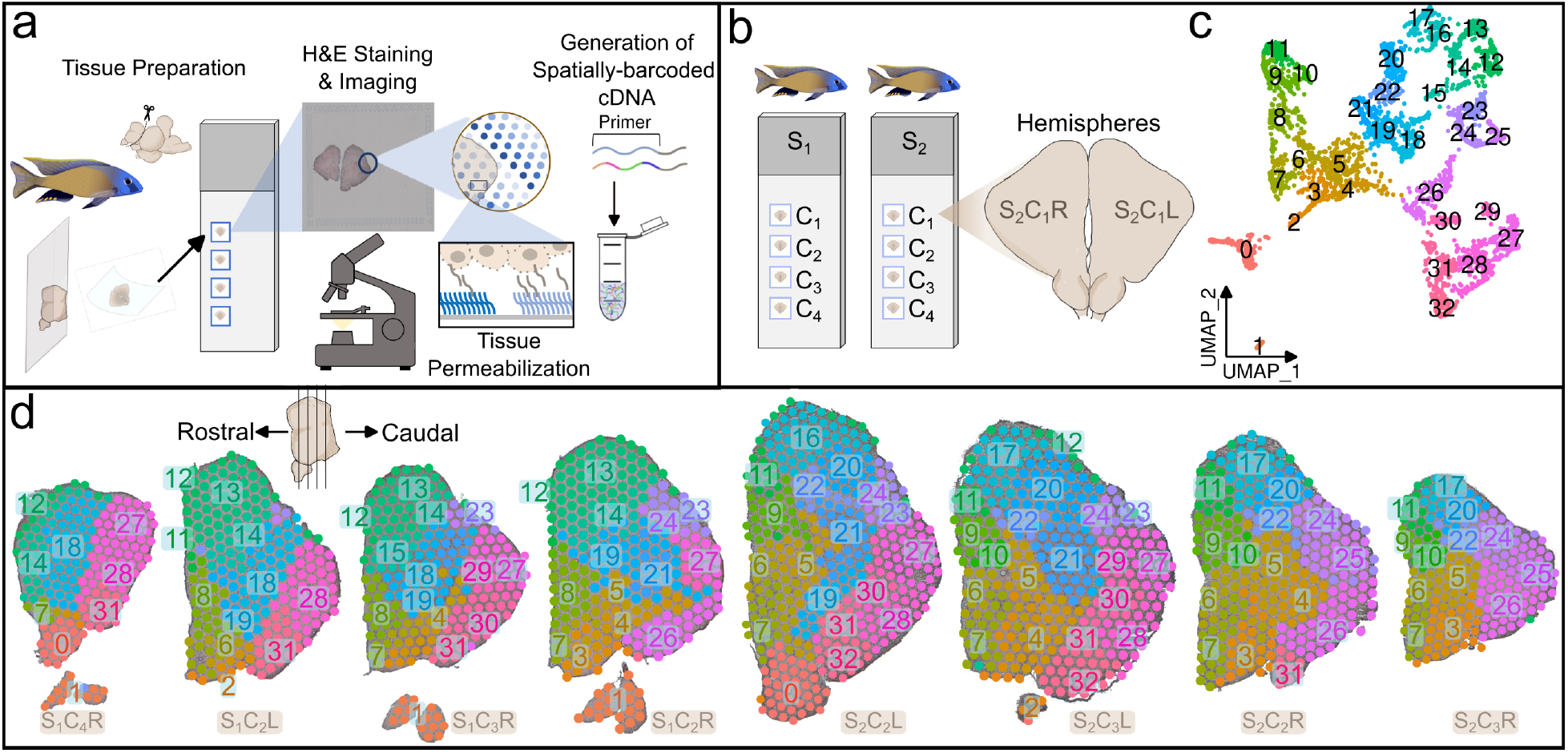
Spatial transcriptomic profiling of the teleost telencephalon. A. Overview of spatial transcriptomics experimental pipeline. 10μm coronal sections from cryopreserved MC telencephala were placed on capture areas of 10X Genomics Visium slides and subsequently fixed, H&E stained, and imaged. Tissue was permeabilized for mRNA capture by barcoded primers on 50μm-diameter spots and spatially-barcoded cDNA was processed for downstream library preparation and sequencing. B. Schematic of experimental design including 4 tissue sections on capture areas (C_1-4_) from two male MC subjects (subject 1, S_1_; subject 2, S_2_), totaling 8 samples processed on Visium platform. Tissue from the left and right telencephalic hemispheres (L and R) were visualized separately. C. Unsupervised gene expression clustering of spots visualized in UMAP space. D. 50μm-diameter spots on select tissue section hemispheres are colored by unbiased clusters from C. The order of tissue sections along rostrocaudal axis was determined through visual inspection of H&E-stained tissue.

### Unbiased clusters represent anatomical regions

Anatomical regions were manually annotated based on tissue cytoarchitecture and expression of select canonical gene markers (Fig. 2A; Supplementary Results) and the distribution of unbiased clusters across these regions was assessed (Fig. S5). We observed high correspondence between anatomical regions and unbiased clusters (Rand index = 0.9487354), suggesting that transcriptional profiles are strongly associated with anatomical context.

**Figure 2.**
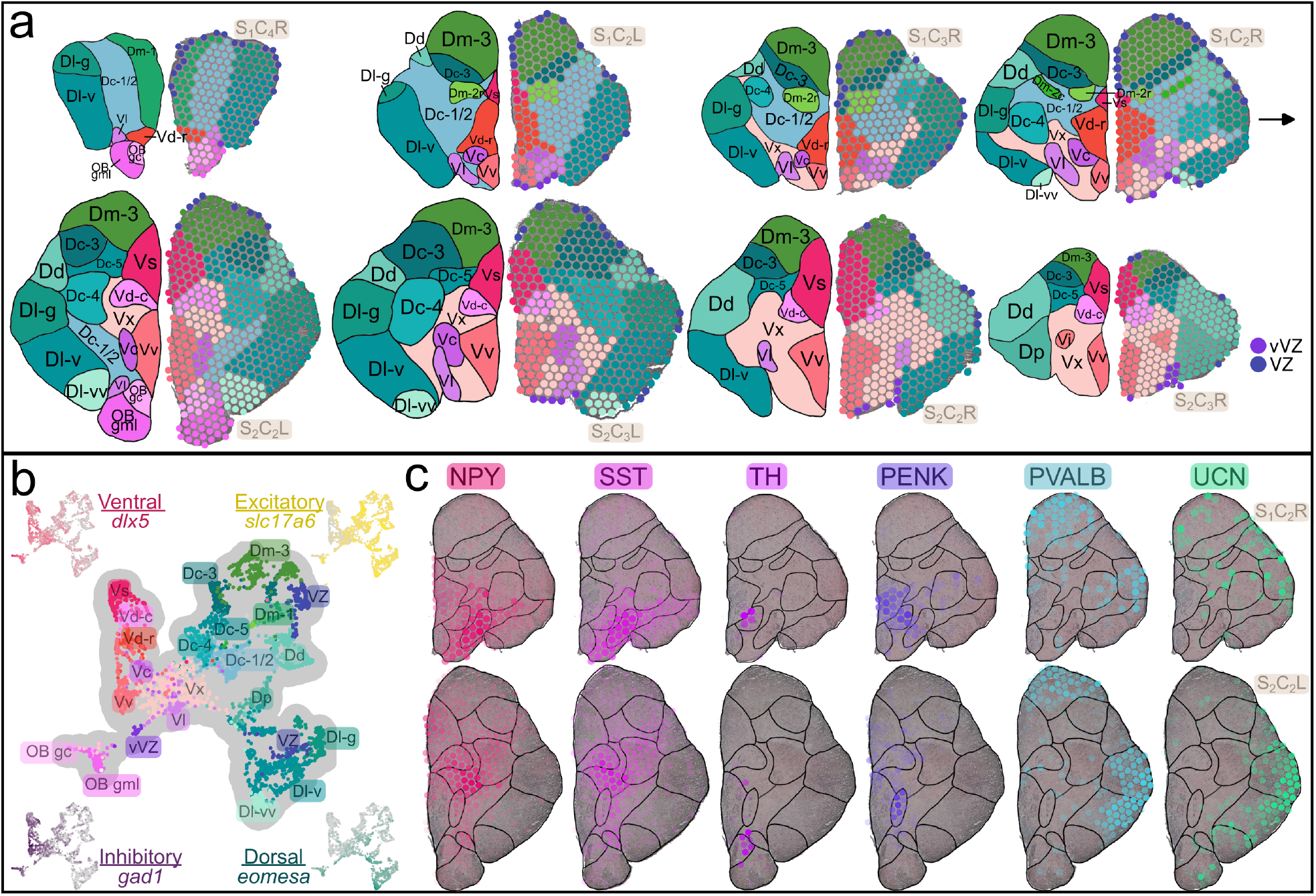
Gene expression patterns in neuroanatomical regions. A. Anatomical annotation of select tissue sections (order same as Fig. 1) with representative coronal atlases shown on the left and spots colored by anatomical region on the right. See Table 1 for abbreviations. B. Spots visualized in UMAP space with anatomical annotations. Also shown is the expression of ventral (*dlx5*), dorsal (*eomesa*), excitatory (*slc17a6*), and inhibitory (*gad1*) marker genes. C. Expression of neuromodulatory genes in representative section hemispheres. Transparency of spots scaled by expression level (spots not expressing the gene are completely transparent).

**Table 1:** Abbreviations.

|  |  |
| --- | --- |
| D | Dorsal (pallial) division of the telencephalon |
| Dc | Central division of D |
| Dc-1,2,3,4,5 | Subdivisions of Dc |
| Dd | Dorsal division of D |
| DI | Lateral division of D |
| DI-d | Dorsal subdivision of DI |
| DI-g | Granular zone of DI |
| DI-v | Ventral subdivision of DI |
| DI-vv | Ventral zone of DI-v |
| Dm | Medial division of D |
| Dm-1,2,3 | Subdivisions of Dm |
| Dm-2c | Caudal part of Dm-2 |
| Dm-2r | Rostral part of Dm-2 |
| Dp | Posterior division of D |
| OB gc | Olfactory bulb granule (internal) cell layer |
| OB gml | Olfactory bulb glomerular/mitral (external) cell layers |
| V | Ventral (subpallial) division of the telencephalon |
| Vc | Central nucleus of V |
| Vd | Dorsal nucleus of V |
| Vd-c | Caudal part of Vd |
| Vd-r | Rostral part of Vd |
| Vi | Intermediate nucleus of V |
| VI | Lateral nucleus of V |
| Vs | Supracommissural nucleus of V |
| Vv | Ventral nucleus of V |
| Vx | Unassigned subdivision of V |
| VZ | Ventricular zone |
| vVZ | Ventral ventricular zone |

The teleost telencephalon is subdivided into pallial (dorsal) and subpallial (ventral) domains which are predominantly populated by glutamatergic or GABAergic neurons, respectively. We identified these divisions in ST data based on expression patterns of conserved pallial (*neurod, eomes*) and subpallial (*dlx2, dlx5*) marker genes (Fig. 2B)^17, 18^. As expected, we observed strong glutamatergic marker gene expression in pallial regions (*slc17a6*, *slc17a7*; Fig. 2B) and GABAergic expression dominating subpallial regions (*gad1*, *gad2*; Fig. 2B)^19^.

Gene expression properties of profiled anatomical regions aligned with patterns previously described in teleost studies, briefly summarized below. Subpallial regions Vv and Vl are considered putatively homologous to the septal formation in mammals^1^. In these regions, we observed expression of markers in agreement with previous reports in fish (Vv: *isl1*, *lhx6, lhx8, and zic1*^17, 20^; Vl: *sst*, *npy* and *crh*^1, 21, 22^; Fig. 2C, Fig. S6). The olfactory bulb granule cell layer (OB gc) expressed dopaminergic cell-type marker tyrosine hydroxylase (*th*)^23, 24^ along with *pax6*, a gene necessary for adult neurogenesis of dopaminergic neurons in the mammalian OB^25^. Vd, Vs, and Vc expressed markers of mammalian striatal medium spiny neurons (MSNs; *penk*, *meis2, six3*; Fig. 2C, Fig. S6)^26, 27^ and genes involved in dopaminergic transmission (*slc18a2*, *slc6a3*; Fig. S6)^23^, in agreement with previously reported similarities between these regions and the mammalian striatum^1, 28^. Teleost pallial regions Dl and Dm are considered the putative homologues of the mammalian medial and ventral pallium, respectively^29^. In zebrafish, expression of *pvalb* has been used to differentiate Dl from the Dm^30^. Interestingly, this pattern is not replicated in our data, as Dm expressed *pvalb7*, albeit at a lower level than in Dl (Fig. 2C; Fig. S6). Urocortin (*ucn*), a member of the corticotropin-releasing factor (CRF) family, was highly expressed throughout the pallium, consistent with previous observations in teleosts (Fig. 2C; Fig. S6)^22, 31^.

The majority of gene expression studies in teleost models have been limited by the necessary identification of gene targets prior to experimentation. The advent of ST technology eliminates this necessity, allowing profiling of thousands of genes simultaneously within tissue organization. In addition to examining previously reported gene expression patterns, we identified novel marker genes demonstrating exclusive expression within telencephalic subregions (Table S4, Fig. S6).

### Anatomical distribution of cell-types in the telencephalon

ST serves as a powerful tool for investigating the expression of thousands of genes within tissue context, but lacks precise cellular resolution as each 50μm-diameter spot may capture RNA from multiple cells and cell-types. Algorithms for cell-type deconvolution of spots address this limitation by estimating the cellular composition of individual spots. We recently profiled the cell-types in the MC telencephalon (38 subjects and 33,674 nuclei) using snRNA-seq^15^. We predicted the anatomical location of these cell-types in our ST data using cell2location^32^(Fig. 3A-B), a top performing tool for cell-type deconvolution of spatial spots^33^. Spots contained on average 8.7±0.086 cells (Fig. S7; Table S5) and most (>92%) were composed of more than one cell-type (4.3±0.032 cell-types per spot). In general, anatomical locations of snRNA-seq cell-types (Fig. 3A-B) agreed with predictions based on previously-reported gene expression patterns in teleosts. For example, cell-types 5.1_GABA and 5.2_GABA, which demonstrated expression of well-known teleost OB marker genes (e.g. *th*, *pax6*)^23–25^, mapped to the OB and 4_GABA cell-types which expressed genes enriched in striatal MSNs (*sp9, six3*) were located in the Vd and Vs regions.

**Figure 3.**
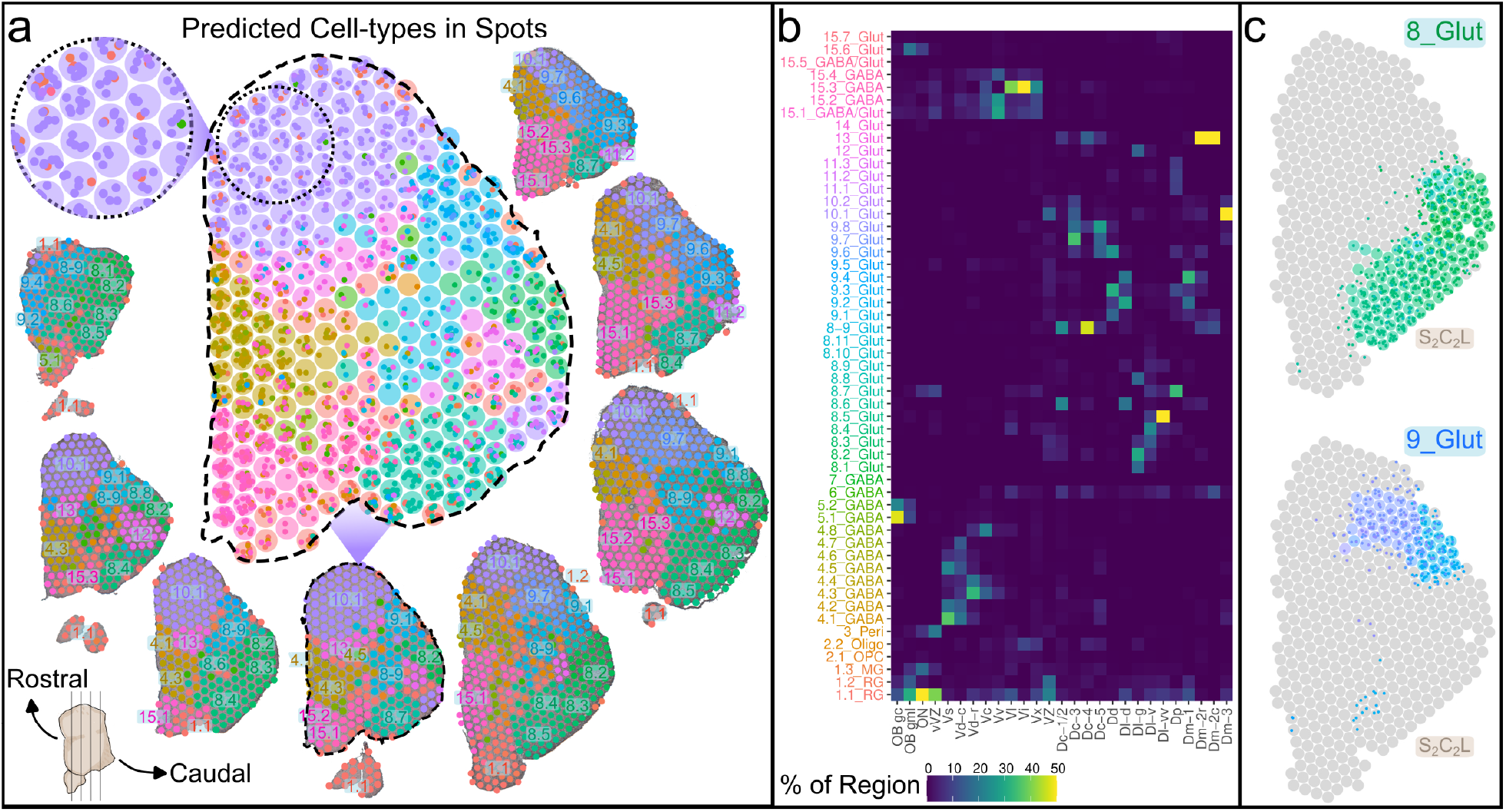
Spatial context of cell-types profiled by snRNA-seq. A. Estimates of snRNA-seq cell-types in ST spots by cell2location. Representative tissue hemispheres are ordered counter-clockwise as in Fig. 1 and spots on the tissue are colored by the cell-type with the greatest number of predicted cells (color of cell-types shown in panel B). Cell-type abundance estimates in a representative tissue hemisphere (S_1_C_2_R) are shown with estimated cells as small dots within spots (larger circles). B. Cell-type composition of annotated anatomical regions (color represents proportion of cell-types composing anatomical regions). C. Anatomical distribution of behaviorally-relevant 8_Glut and 9_Glut cell-types in S_2_C_2_L.

Next, we investigated the spatial locations of cell-types implicated in bower-building, a courtship behavior wherein male MCs build structures by spatially manipulating sand to attract a mate^34^. Converging lines of evidence from our snRNA-seq experimental assays supported involvement of pallial cell-types 8_Glut and 9_Glut in bower-building. These cell-types exhibited behavior-associated changes in gene expression and neurogenesis. Additionally, 9_Glut was the only cell-type to show significantly elevated building-associated expression of immediate early genes, which are indicators of neuronal excitation. 8_Glut cell-types were predicted to span the Dl and Dp while 9_Glut was mainly located in the Dd (Fig. 3C). Furthermore, the spatial locations of non-neuronal cell-types were identified. For example, subpopulations of radial glia were located along the ventricular zone in areas bordering behaviorally-relevant populations, indicating a potential supply of new neurons to these regions^15^. Our ST data of the *Mchenga conophoros* (MC) telencephalon is a valuable resource for identifying the anatomical location of genes and cell-types in MC and any closely related cichlid species. Furthermore, by resolving the spatial distribution of cell-types, we gain a more comprehensive atlas of the cichlid telencephalon and can further analyze these cell-types with added spatial context.

### Conserved telencephalic cell-types in teleosts and mammals

Around 350-450 million years have passed since the last common ancestor of teleost fish and mammals^35^. Despite a vast array of studies aimed at disentangling the putative mammalian homologs of anatomical regions in fish, obstacles such as the ‘everted’ teleost pallium have made this challenging. To investigate the evolution of cell-types within the telencephalon, we analyzed conserved transcriptional signatures present in comparable cichlid and mouse forebrain datasets. First, we evaluated two common methods for comparing single-cell RNA-seq (scRNA-seq) data from distantly related species: correlations of expression of common marker genes and SAMap^13^, which uses sequence similarity of proteins from species being compared as weights while integrating their datasets together. Each method was used to compare a mouse scRNA-seq dataset^36^ to a downsampled version of itself and the matrix of cell-type similarity scores produced by each method was compared to the identity matrix. SAMap had the greatest coefficient of determination (SAMap R^2^=0.436; Fig. 4A) and was therefore primarily used to compare cichlid and mouse^36^ telencephalic cell-types (Fig. S8). Despite vast evolutionary distances, sequence similarity-aware integration of these datasets projected nuclei/cells together in UMAP space (Fig. 4B). Next, a similarity score was calculated based on the mean knn neighbors between cichlid and mouse cell-types and the score was compared to permutations (Fig. 4C; Table S6). Cell-types with similarity scores greater than all permutations (q_perm_=0; n_perm_=1,000) are outlined below.

**Figure 4.**
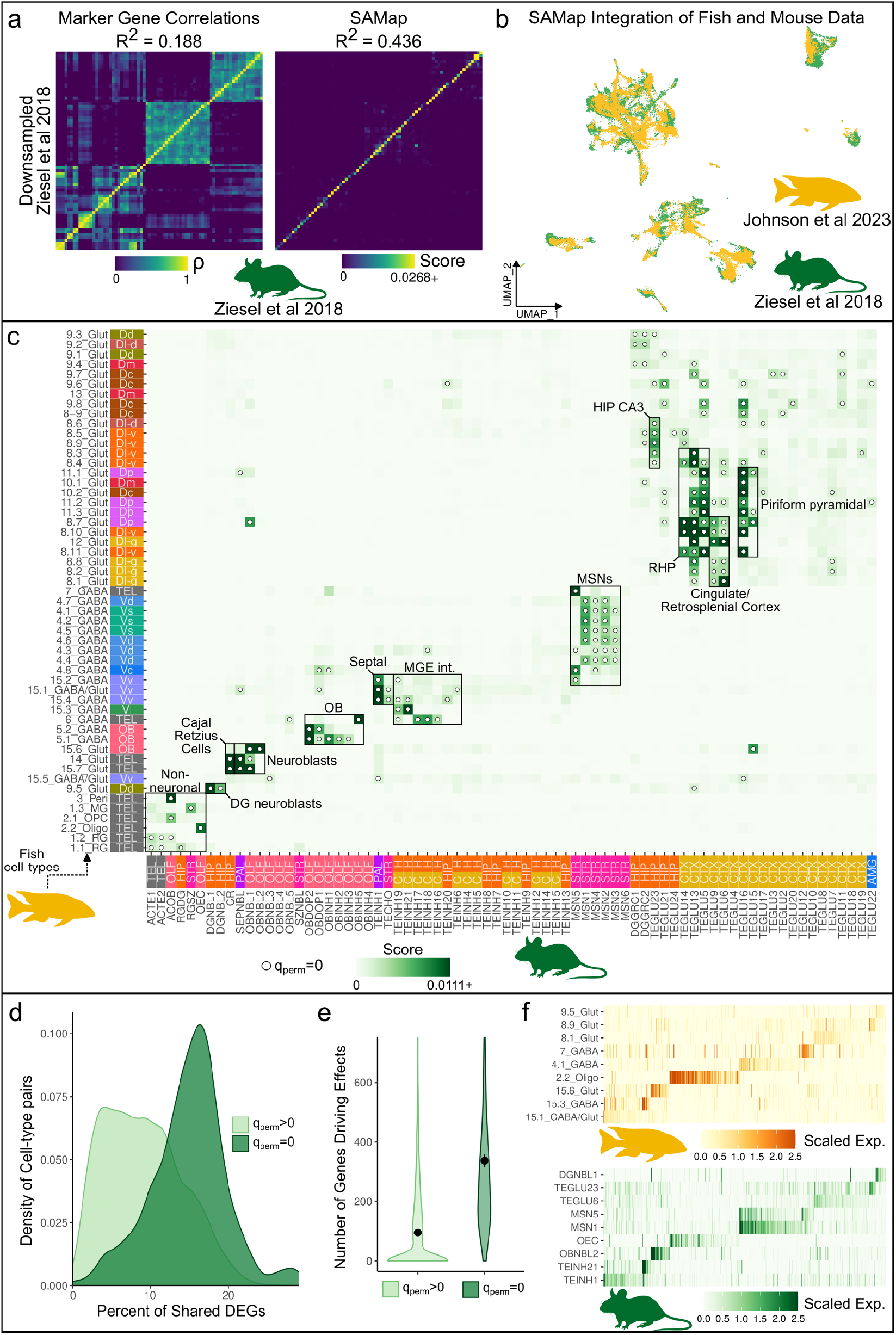
Comparative analysis of cell-types in teleosts and mice. A. Comparison of common methods for comparing distantly related scRNA-seq datasets: correlations of expression of marker genes and SAMap. To evaluate the accuracy of these methods, they were used to compare a mouse telencephalon dataset to a downsampled version of itself. The output of these methods are shown with cell-types from the downsampled dataset on the y-axis and cell-types from the full dataset on the x-axis. The accuracy of the outputs were found by calculating the R^2^ with the identity matrix. SAMap had the greatest accuracy (R^2^=0.436). B SAMap projects cichlid nuclei (n=33,674) and mouse cells (n=59,492) from telencephalon together in UMAP space. Cichlid nuclei are ordered to the front. C. The mean number of k-nearest-neighbors, computed by SAMap, between cichlid and mouse cell-types were used as a similarity score and were compared to permutations. Cichlid cell-types on the y-axis with color labels for their predicted spatial location. Mouse cell-types on the x-axis with color labels for their annotated region of origin^36^: AMG=amygdala, CTX=cortex, OLF=olfactory, HIP=hippocampus, PAL=pallidum, STR=striatum, TEL=telencephalon. Darker green represents greater transcriptional similarity and dots represent similarity scores greater than all permutations (q_perm_=0). D. Conserved marker genes (e.g. differentially expressed genes) between cichlid and mouse cell-types were found. Distribution of the percent conserved markers for significant (q_perm_=0, dark green) and not-significant (q_perm_>0, light green) cell-type pairs. Cell-type pairs with significant similarities had significantly more conserved marker genes than other cell-type pairs (Welch Two Sample t-test; p=4.843308e-40, t=16.54035). E. Violin plot of the number of genes driving effects for cell-type pairs with significant and non-significant cell-type pairs. Dots indicate the mean value and vertical lines represent the standard error. Possible outliers (<1^st^ or >99^th^ percentile) are not visualized. Cell-type pairs with significant similarities also had significantly more genes driving effects than other cell-type pairs (Welch Two Sample t-test; p=1.151187e-22, t=11.1492). F. Scaled expression of DEGs driving select cell-types relationships (n_genes_=652).

We found strong cross-species correspondence of major cell-type classes, with glutamatergic, GABAergic, and non-neuronal populations demonstrating significant transcriptional similarities between cichlid and mouse. Cichlid *Plp1+* oligodendrocytes (2.2_Oligo) were significantly similar to murine olfactory ensheathing cells (OEC) and *Fabp7+* radial glia populations (1.1_RG and 1.2_RG) resembled mouse telencephalon astrocyte cell-types (ACTE1-2). In the teleost telencephalon, radial glia line the pallial and subpallial ventricular zones (VZ), which resemble the main neurogenic niches of the adult mammalian brain, the dentate gyrus and subventricular zone^37^. Radial glia function as stem cells, giving rise to new neurons and glia throughout adulthood^37^ and are considered astroglial due to their expression of classical astrocyte markers^38^ despite lacking the stellate morphology characteristic of mammalian astrocytes. Additionally, the teleost pallial VZ contains a population of non-glial proliferative progenitors largely composed of neuroblasts^39^. Cichlid 9.5_Glut, concentrated along the pallial VZ, expressed markers of neuroblasts (*sox4*, *sox11*, *mex3a* and *zeb2*) and bore molecular similarity to *Sox4*+ granule neuroblasts of the hippocampal dentate gyrus (DGNBL1-2). The most abundant neuroblast population in the teleost telencephalon forms a rostral migratory stream-like structure in the subpallial VZ, migrating into the OB^37^ similar to observations in the subventricular zone of rodents^40^. Indeed, the cichlid cell-type located in the glomerular/mitral layer (gml) of the OB (*Eomes+/Tbx21+* 15.6_Glut) resembled mouse glutamatergic OB neuroblast populations (OBNBL1-2). Cichlid *Tp73+* 15.7_Glut and *Cacna2d2+* 14_Glut, which were similar to several mouse neuroblast populations, additionally resembled dentate gyrus Cajal-Retzius cells (CR). In the mouse brain, these cells play critical roles in hippocampal and neocortical development and radial glia migration^41^.

In the mammalian brain, GABAergic lineages are derived largely from the ganglionic eminence progenitor domains (e.g. MGE, LGE), and early subpallial territories in the embryonic teleost brain have been proposed as MGE/pallidal-like or LGE/striatal-like based on regionalized expression of neuroregulatory genes^42^. Several cichlid GABAergic cell-types were significantly similar to mouse LGE-derived populations, including *Six3+* 4_GABA cell-types in the Vd and Vs with mouse striatal MSNs (MSN1-4). A recent cross-species comparison between the zebrafish and mouse forebrain also found strong correspondence between mammalian MSNs and zebrafish subpallial cell-types expected to be located near the Vd^12^. LGE-derived *Th*+ cell-types located in the cichlid and mouse OB mapped to each other with great specificity (5.1_GABA and 5.2_GABA to OBDOP). We observed strong molecular conservation between cichlid *Sst, Npy+* 15.3_GABA, concentrated in the Vl, with mouse Sst-class interneurons (TEINH21, TEINH19) and *Pvalb+*/*Nxph1+* 6_GABA with mouse interneurons in the hippocampus and cortex (TEINH17-18). Finally, several cichlid *Isl1*+ cell-types located in the Vv (15.1_GABA/Glut, 15.2_GABA and 15.4_GABA) were similar to a septal cell-type in the pallidum (*Lhx6+* TEINH1), with 15.1_GABA/Glut mapping to additional mouse septal cell-types (TEHCO, cholinergic interneurons; SEPNBL, septal neuroblasts).

Among glutamatergic cell-types, *Bhlhe22*+ 8_Glut cell-types located in the ventral-most Dl (Dl-vv; 8.3_Glut, 8.5_Glut, 8.6_Glut, and 8.9_Glut) were transcriptionally similar to a mouse hippocampal CA3 cell-type (TEGLU23). We also observed similarity between several *Cck*+ 8_Glut cell-types in the Dl-v and mouse cell-types of the subiculum (TEGLU13-14), a mammalian structure in the retrohippocampal formation that mediates hippocampal-cortical communication^43^. Notably, several cichlid populations were transcriptionally similar to cortical projecting excitatory neurons, including 8.1_Glut and 12_Glut, both located in the Dl-g, with mouse neocortical cell-types from cingulate/retrosplenial areas (TEGLU6, TEGLU9). *Emx3*+ populations which mapped to the Dm (10_Glut) and Dp (11_Glut and 8.7_Glut) bore similarity to a mouse cell-type in cortical entorhinal superficial layers (TEGLU5) as well as to piriform pyramidal cells (TEGLU15-17).

To robustly determine molecular signatures of conserved cell-types found by SAMap, we used two methods: conserved marker genes (DEGs in common between cell-types) and genes found by SAMap that drive relationships. Both methods require that each gene is differentially expressed, however the thresholds used are different. The former is more strict (see Methods) and is orthogonal to SAMap. Cichlid-mouse cell-type pairs exhibiting significant similarity had a mean of 57.57812±1.932748 conserved marker genes and a mean of 336.9583±21.57531 genes driving effects reported by SAMap (Table S7), both of which were significantly greater than in non-significant cell-type pairs (Fig. 4D-E). Expression of genes driving select significant cell-type pairs is shown in Figure 4F. Additionally, to determine the robustness of these results, a comparison to an additional mouse telencephalon dataset^44^ was performed and consistent similarities between cichlid and mouse cell-types were observed (Supplementary Results; Fig. S9). In summary, our results suggest molecular conservation of non-neuronal populations, neuroblasts, olfactory cell-types, MSNs, hippocampal, and cortical cell-types.

### Conserved telencephalic cell-types in teleosts and tetrapods

Across vertebrate lineages, the telencephalon demonstrates both commonalities shared between taxa as well as marked, specialized differences and many hypothesized brain region homologies in non-mammalian vertebrates are unclear and actively debated. To further investigate conserved telencephalic populations in vertebrates, we performed cross-species comparisons between cichlid cell-types and datasets from tetrapod forebrain regions (Fig. 5A-C; Table S8), including amphibians (axolotl telencephalon^9^) and sauropsids (turtle pallium^7^, songbird HVC and RA pallial regions^8^). Following the same approach previously described, SAMap was used to integrate cichlid nuclei with datasets from each species separately, revealing several consistent patterns outlined below.

**Figure 5.**
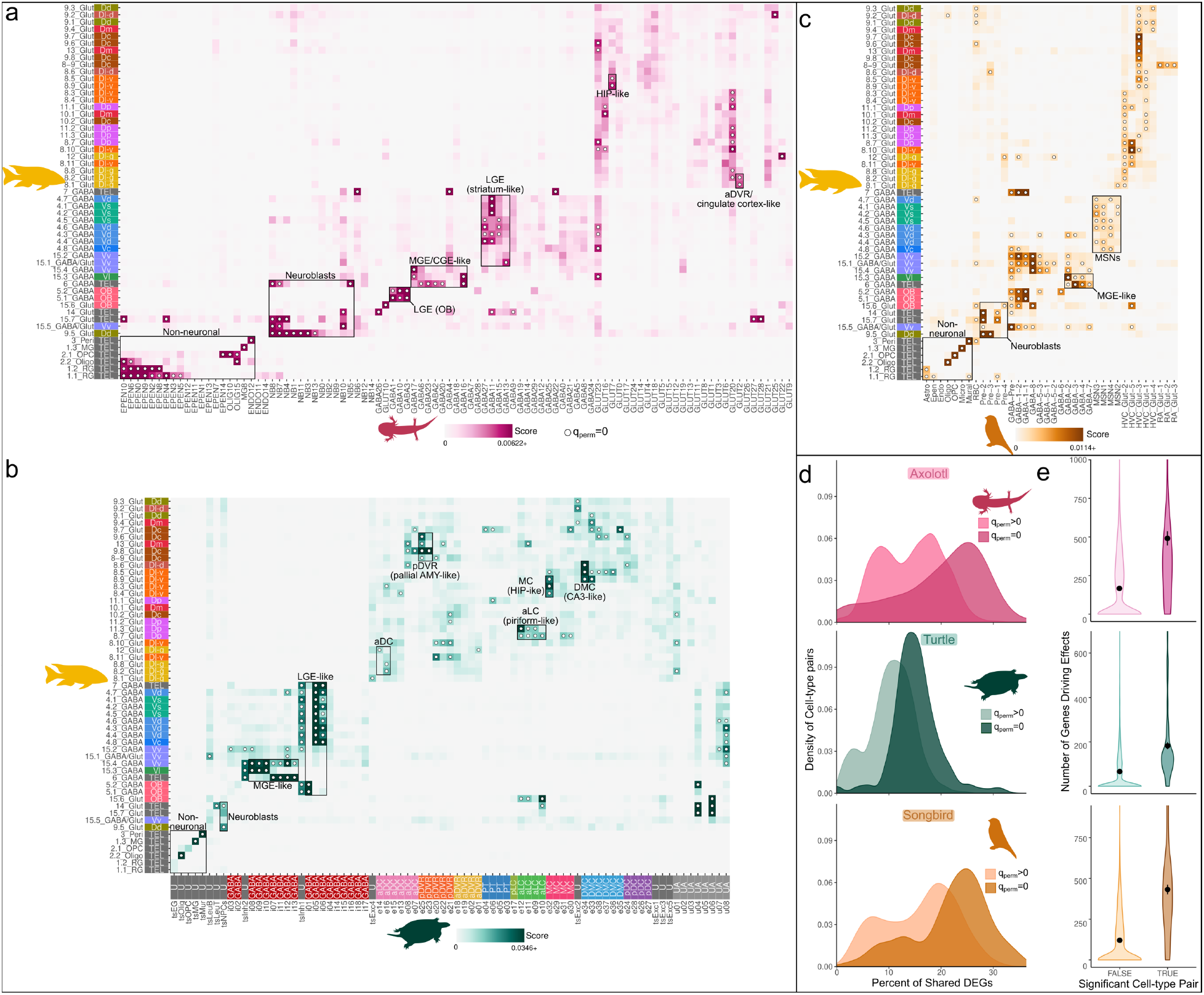
Comparative analysis of cell-types in cichlids to axolotls, turtles and songbirds. A-C. Pairwise transcriptional comparisons of cichlid telencephalic cell-types to axolotl (A), turtle (B) and songbird (C) cell-types using SAMap. Cell-type pairs with higher similarity scores indicate greater transcriptional similarity and dots denote similarity scores greater than all permutations. Turtle cell-types are colored by their inferred region of origin with the following additional labels: U=unknown, UA=unassigned. D. Distribution of the percent of shared DEGs between significant (q_perm_=0) and not-significant (q_perm_>0) cell-type pairs. Cell-types pairs with significant similarities between species have more DEGs in common than other cell-types pairs (Welch Two Sample t-test; axolotl p=2.170296e-10; turtle p=5.540887e-25; songbird p=1.630375e-19). E. Violin plot of the number of genes driving effects for cell-type pairs with significant and non-significant cell-type pairs. Dots indicate the mean value and vertical lines represent the standard error. Possible outliers (<1^st^ or >99^th^ percentile) are not visualized. Cell-type pairs with significant similarities also had significantly more genes driving effects than other cell-type pairs (Welch Two Sample; axolotl: p=1.792532e-10, t=7.076042; turtle: p=1.687547e-11, t=7.274479; songbird: p=5.675997e-20, t=10.48941).

Our comparisons supported correspondence of major non-neuronal cell-types, including cichlid astroglia-like radial glia cell-types to axolotl ependymoglia (*Slc1a2+*) and songbird astrocytes (*Fabp7+*). Cichlid microglia (1.3_MG; *Csf1r+*) and oligodendrocytes (2.2_Oligo; *Plp1+*) consistently mapped to their counterparts in all vertebrate comparisons and oligodendrocyte precursor cells (2.1_OPC, *Olig2+*) showed one-to-one mapping in songbirds. As with the comparison to the mouse telencephalon, *Sox4+* 9.5_Glut demonstrated significant similarity to neuroblasts in all comparisons. Taken together, these findings point towards strongly conserved molecular signatures present in critical non-neuronal populations across vertebrates.

Cichlid GABAergic cell-types demonstrated consistent patterns across comparisons with other vertebrates. Cichlid OB GABAergic cell-types (*Trh+* 5.1_GABA and *Th+* 5.2_GABA) resembled olfactory LGE-derived cell-types (axolotl GABA1, GABA3 and GABA10; turtle i01) and songbird LGE-derived GABA-1-1 and GABA-1-2. Cichlid *Six3+* 4_GABA cell-types similar to mouse striatal MSNs also resembled songbird MSNs from Area X and LGE-derived cell-types in axolotl (GABA11) and turtle (i04-i06). In line with results from our cichlid-mouse analysis, we observed strong transcriptional similarity between vertebrate MGE-derived populations and cichlid *Pvalb6*+/*Nxph1+* 6_GABA (axolotl GABA2 and GABA6; turtle i07, i11-i13; and songbird GABA-3 and GABA-4) and *Sst+/Npy+* 15.3_GABA (axolotl GABA17; turtle i08-i10; songbird GABA-2).

Additionally, we found evidence of conserved molecular features in several glutamatergic cell-types. Similar to our cichlid-mouse comparison, a subset of Dl-v 8_Glut cell-types (8.5_Glut, 8.6_Glut and 8.9_Glut) consistently grouped with hippocampal-like cell-types in vertebrate datasets. These populations resembled turtle dorsal medial cortex cell-types (DMC; e33-e38), determined as transcriptionally similar to mammalian hippocampal CA populations^7^, and HVC_Glut-3 in songbirds, which previous analysis found resembled the mammalian hippocampus and deep cortical layers^8^. Additionally, we found similarity between 8.5_Glut and 8.9_Glut with an axolotl cell-type (GLUT7) reported as hippocampal^9^. Cichlid Dm-2r (13_Glut) and Dc (9.6_Glut, 9.8_Glut) cell-types resembled a turtle population in the posterior dorsal ventricular ridge (pDVR; e25), reported as similar to the mammalian pallial amygdala^7^. Cell-types located in the *Emx1+* Dp (8.7_Glut, 11.3_Glut) resembled a subset of turtle cell-types in the anterior lateral cortex (aLC; e09-e12), the putative reptilian homologue of the piriform cortex^7^. Additionally, 8.7_Glut mapped to a songbird cell-type (HVC_Glut-5) determined by previous analysis to resemble mammalian cortical populations, including the piriform cortex^8^. Cichlid Dl-g cell-types were similar to populations in the turtle aDC (8.2_Glut and 8.8_Glut with e16; 12_Glut with e14) and aDVR (12_Glut with e01), as well as to songbird and axolotl cell-types determined as similar to the turtle aDVR (12_Glut with songbird HVC_Glut-2 and HVC_Glut-5; 8.1_Glut with axolotl GLUT2)^8, 9^. Axolotl GLUT2 was derived mainly from microdissections of the telencephalon containing the ventral pallium^9^, a region found by independent analysis to contain populations similar to the turtle aDVR^10^.

Our analyses revealed molecular conservation between cell-types in the cichlid telencephalon and those in the forebrains of multiple vertebrate lineages. In support of these findings, cell-types with significant similarities shared more marker genes and were driven by a greater number of genes than others (Fig. 5D-E; Table S9). For populations exhibiting the most striking conservation across species, we identified the core set of genes defining these evolutionary conserved cell-types (Fig. S10-12; Table S10).

### Transcriptional similarities of anatomical regions in teleosts and tetrapods

Our atlas of the cichlid telencephalon provides a powerful tool to explore transcriptional similarities of telencephalic regions across species, amidst competing hypotheses about putative mammalian homologies of teleost brain regions. Using SAMap, we integrated ST cichlid data with anatomically-annotated scRNA-seq data from turtle pallial microdissections and separately with ST data from the mouse brain^45^. Below we outline significant relationships between brain regions supported by and consistent with cell-type relationships previously described.

Comparison of cichlid-turtle pallial brain regions revealed the following significant similarities between: Dl-vv and the DMC, Dp and the aLC, Dm-2r and the pDVR, and the Dl-g and the aDVR (Fig. 6A; Table S12). The relationship between the Dl-vv and the DMC was supported by our cell-type analysis (Fig. 5B) which found that cell-types mapping to these regions (cichlid 8.5_Glut and turtle e34) resembled the mammalian CA3, reported here and by Tosches et al^7^. Similarly, the reptilian aLC and cichlid Dp are both reported as putative homologues of the mammalian olfactory (piriform) cortex^7, 43^ and their homology to each other was further supported by our cell-type comparison (cichlid 8.7_Glut and 11.3_Glut with turtle e09-e12; Fig. 5B). Cichlid cell-types located in the ventromedial Dm (Dm-2r, Dm-2c) aligned with turtle pDVR cell-types (13_Glut with e25; Fig. 5B), both regions which have been compared to the mammalian pallial amygdala ^7, 46^.

**Figure 6.**
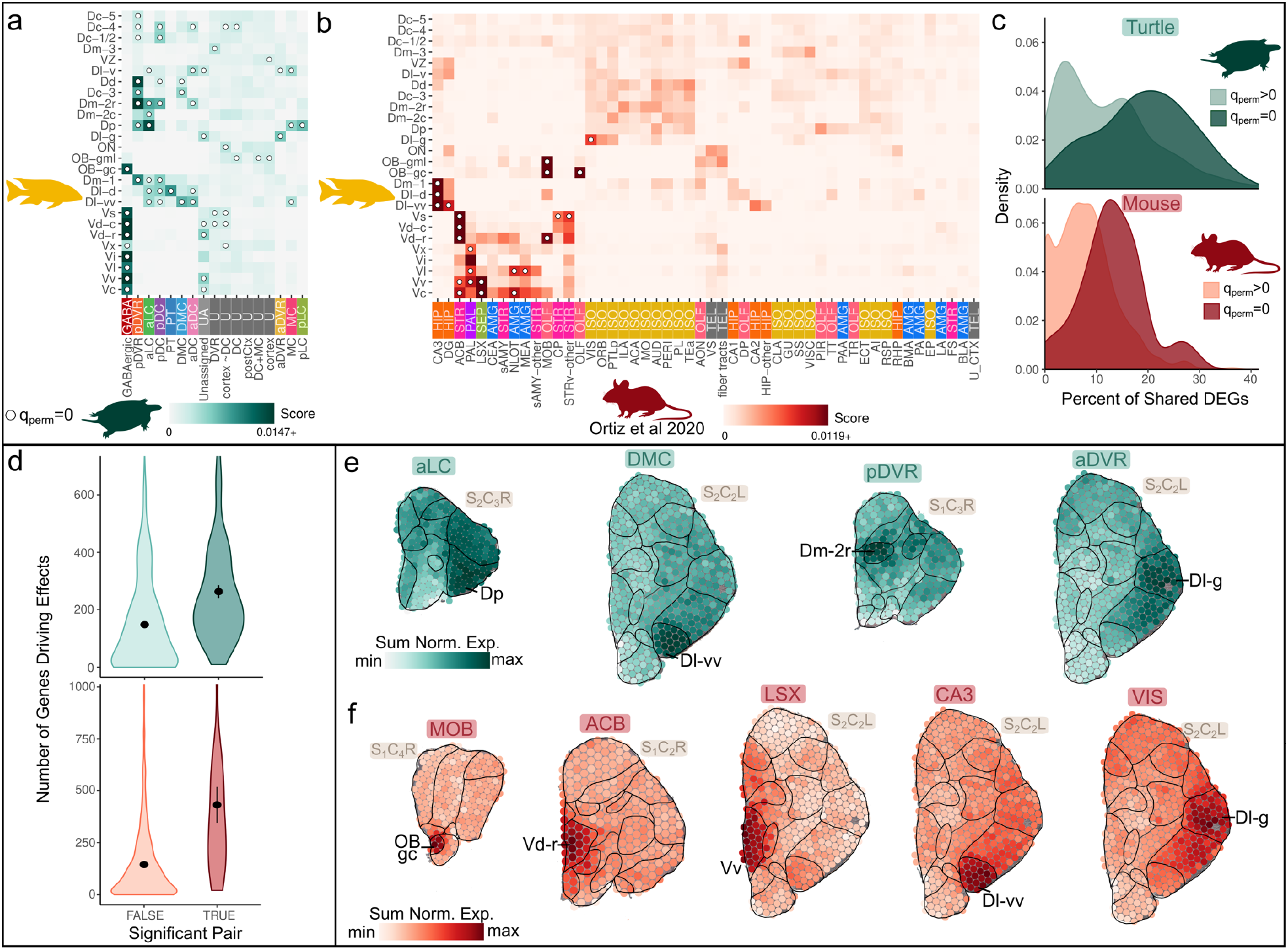
Comparison of neuroanatomical regions in cichlids to turtles and mice. A. Transcriptional comparisons of annotated cichlid telencephalic ST anatomical regions to anatomical annotation of turtle cells using SAMap. Greater similarity scores indicate greater transcriptional similarity and dots denote significance. B. Transcriptional comparisons annotated cichlid telencephalic ST anatomical regions to ABA region annotations of mouse ST data using SAMap. AMG=Amygdala, HIP=Hippocampal formation, ISO=Isocortex, OLF=Olfactory areas, STR=Striatum, PAL=Pallidu, SEP=Septum and TEL=Telencephalon. C. Distribution of the percent of shared DEGs between significant (q_perm_=0) and not-significant (q_perm_>0) anatomical regions. Regions with significant similarities have more DEGs in common than other cell-types (turtle p=2.390385e-07, t=5.678015; mouse p=4.940415e-04, t=4.032538). D. Violin plot of the number of genes driving effects for cell-type pairs with significant and non-significant cell-type pairs. Dots indicate the mean value and vertical lines represent the standard error. Possible outliers (<1^st^ or >99^th^ percentile) are not visualized. Cell-type pairs with significant similarities also had significantly more genes driving effects than other cell-type pairs (Welch Two Sample; turtle: p=4.868006e-06, t=4.886118; mouse: p=0.003745793,t=3.324379). E. Expression of genes driving select significant relationships between cichlid and turtle anatomical regions (turtle aLC and cichlid Dp; turtle DMC and cichlid Dl-vv). F. Expression of genes driving select significant relationships between cichlid and mouse anatomical regions (mouse MOB and cichlid OB gc; mouse ACB and cichlid Vd-r; mouse LSX and cichlid Vv; mouse CA3 and cichlid Dl-vv; mouse VIS and cichlid Dl-g).

Consistent with previous studies in fish, our cichlid-mouse comparison of subpallial regions revealed strong correlation between the following regions: cichlid OB gc and OB gml with the main olfactory bulb (MOB), the Vd with the striatum (Vd-r and Vd-c with nucleus accumbens, ACB), and the Vv with the lateral septal complex (LSX) (Fig. 6B; Table S11). Results from this analysis also support the proposed homology between the Dp and the mammalian piriform cortex, in agreement with the above comparison to turtle brain regions and with the partial eversion model of the teleost telencephalon^2^. The Dl-v is considered the most-likely homologue of the vertebrate hippocampal pallium^5, 43, 47, 48^, and we found that a dense subpopulation at its ventral-most pole, the Dl-vv, bore strong similarity to the hippocampus. The Dl-g has been previously proposed as the putative homologue of the hippocampal DG based on its exclusive expression of *prox1* in zebrafish histochemical studies^18^, but this gene was not strongly expressed in cichlid pallial cell-types. Instead, our analysis suggests that the Dl-g shares greater transcriptional similarities with visual areas of the neocortex (VIS), in agreement with neocortical signatures observed in cell-types composing Dl-g (8.1_Glut and 12_Glut; Fig. 4).

Again, we found that populations demonstrating significant transcriptional similarity shared significantly more marker genes and were driven by a greater number of genes than others (Fig. 6C-D; Table S12). The molecular similarity of these regions was supported by the sum of the normalized expression of their shared marker genes (Fig. 6E-F).

### Anatomical subregions in vertebrates exhibiting transcriptional similarity to the mammalian neocortex

The teleost brain lacks a layered cortex and telencephalic populations that resemble those present in this elaborated mammalian structure remain a matter of debate. A popular theory is that no such homologue exists, while others propose the Dc^2, 30^ or the Dl-d/g region as a phenotypic homologue of the dorsal pallium or neocortex^2, 3, 49^. In both of our cichlid-mouse comparisons, Dl-g and its predominant cell-types were transcriptionally similar to neocortical populations. Cell-type comparisons to the turtle pallium revealed similarities between cell-types in the Dl-g (12_Glut) and turtle aDC (e14), which previous analyses found resembled to the neocortex when restricted to transcription factors^7^. Additionally, the Dl-g and the same Dl-g cell-type (12_Glut) were transcriptionally similar to the turtle aDVR and one of its cell-types (e01). Previous analysis of the turtle aDVR using effector genes and transcription factors found strong correlations to the neocortex, but not when analyzing only transcription factors^7^. To gain further insight on the relationship of these structures, we used SAMap to compare turtle and mouse scRNA-seq datasets.

This analysis supported significant similarities between aDC cell-types (including e14) and an aDVR cell-type (e01) to the neocortex (Fig. 7A; Fig. S13). The composition of genes driving these relationships by transcription factors, neuromodulatory ligands and receptors did not significantly differ between the aDC with the neocortex, aDVR with the neocortex, and other significant cell-type pairs (Fig. 7B; χ^2^ test, p_tf_=0.4485192, t_tf_=1.6036076; p_ligand_=0.1386000, t_ligand_=3.9523268; p_receptor_=0.7211721, t_receptor_=0.6537549; Table S13). Similarly, genes driving effects between cichlid Dl-g and mammalian neocortex cell-types did not differ in composition compared to other significant cichlid-mouse cell-type pairs (χ2 test, p_tf_=0.5225745, t_tf_=0.408809196; p_ligand_=0.9383825, t_ligand_=0.005975749; p_receptor_=0.2074504, t_receptor_=1.589130627). Additionally, genes driving effects between cichlid Dl-g and turtle aDC, cichlid Dl-g and turtle aDVR, and other significant cell-type pairs differed only in composition of neuromodulatory ligands (χ2 test, p_tf_=0.47667411, t_tf_=1.481844; p_ligand_=0.02823909, t_ligand_=7.134096; p_receptor_=0.38646811, t_receptor_=1.901412). Similar observations were made for the composition of conserved marker genes (Supplementary Results; Fig. S14)

**Figure 7.**
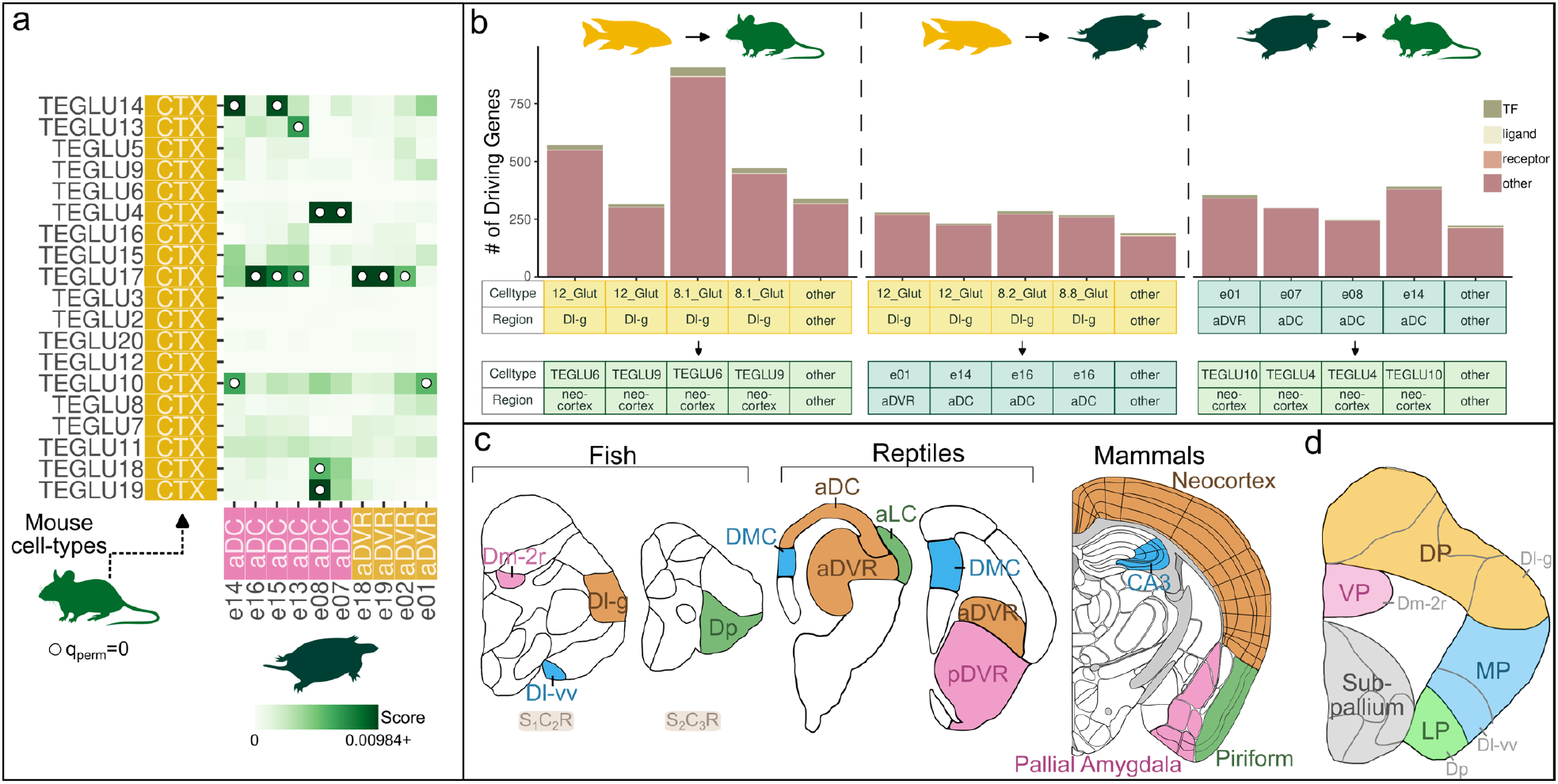
Neocortical signatures in reptiles and fish. A. Transcriptional comparison of cell-types in the turtle and mouse forebrain using SAMap. Results for cell-types from the turtle aDC and the aDVR to cortical mouse cell-types are shown and the full results can be found in the Supplement. B. Composition of genes driving significant cichlid-mouse, cichlid-turtle and turtle-mouse cell-type pairs by transcription factors, neuromodulatory ligands and receptors. No significant differences in composition were found. C. Representative schematic of coronal sections reflecting proposed relationships between telencephalic brain regions in fish, reptiles (modified from Tosches *et al.* 2018^7^) and mammals^45^. D. Representative coronal schematic demonstrating teleostean equivalents of pallial divisions based on the partial eversion model and supported by our results. Colors represent the teleostean equivalents of the dorsal, medial, lateral, and ventral pallium divisions (separated by solid lines). Yellow=dorsal pallium, pink=ventral pallium, blue=medial pallium, green=lateral pallium and gray=subpallium. Gray lines represent subregions of the teleost telencephalon.

Based on our findings and previous literature, we propose the following brain regions contain molecularly conserved populations in cichlids, turtles and mice: Dl-g, aDVR/aDC and neocortex; Dl-vv, DMC and CA3; Dm-2r, pDVR and pallial amygdala; Dp, aLC and piriform cortex (Fig. 7C). These findings support the putative pallial divisions outlined in the partial eversion model, which proposes the Dl-v as medial pallial, the ventromedial Dm as ventral pallial, the Dp as lateral pallial, and the dorsal Dl as dorsal pallial^2^ (Fig. 7D).

## DISCUSSION

Here we create a spatially resolved-molecular atlas of teleost cichlid telencephalon, harnessing the benefits of snRNA-seq and ST to investigate cell-types sequenced at single cell resolution within tissue architecture. We find remarkable correspondence between transcriptional profiling and neuroanatomy, whereby both ST clusters and snRNA-seq cell-types align strongly with well-described neuroanatomical subregions of the fish telencephalon. Comparable observations have been made in the mouse brain: unbiased clusters of ST data recapitulated isocortical layer divisions^45^ and cell-type deconvolution tools of ST data have been shown to accurately determine the position of cell-types with well-characterized spatial locations^32^. Using a comparative approach that considers protein sequence similarity, we identify a suite of non-neuronal and neuronal populations that show strongly conserved patterns of gene expression across all major vertebrate lineages.

We find strong evidence of cichlid GABAergic populations that closely resemble mammalian MGE-derived *Sst+* and *Pvalb+* interneuron classes and LGE-like populations, including OB cell-types and striatal MSNs. Cichlid MSN-like cell-types are concentrated in the Vd, a region which receives substantial dopaminergic input and is widely considered homologous to the mammalian striatum^24, 28^. Our results also support a previously proposed homology between the cholinergic Vv and the tetrapod septum^50, 51^. Various studies have shown that both the striatum and septum, and their putative homologues in fish, play important roles in mediating evolutionarily-relevant social behaviors^51–54^. Our results strongly suggest that core cell-types in these subpallial regions are conserved across vertebrates, perhaps due to essential behavioral functions.

The pallium has presented perhaps the most challenging puzzle in vertebrate brain evolution due to its markedly unique adaptations across vertebrate clades. The teleost telencephalon exhibits an everted morphology, but the extent of eversion is unclear. Some models suggest a complete eversion of all pallial zones, while others propose only a partial eversion^2–6^. Consequently, these models differ in the teleostean equivalents of mammalian dorsal, lateral, and ventral pallium divisions, but agree that the medial (hippocampal) pallium in fish is displaced in a dorsolateral location^43^. The hippocampal formation, generally considered highly conserved across vertebrates^55^, functions in memory, spatial navigation and learning^56, 57^, and behavioral studies in goldfish demonstrate that the ventral subdivision of the dorsolateral pallium (Dl-v) is essential for these tasks^48, 58, 59^. Our results further support this homology, as the ventral-most Dl-v population exhibits striking similarities to the mammalian hippocampal CA3 region and hippocampal populations in multiple vertebrate species. These similarities are particularly interesting given recent findings that highlight a role for this region in bower-building, a social courtship behavior involving the spatial manipulation of sand^15^.

Disagreement between eversion models is rooted partly in the caudolateral position of the Dp, which is the main pallial recipient of secondary olfactory input in the fish telencephalon^43^ and thus the most likely homologue of the mammalian lateral (olfactory) pallium. Furthermore, studies in zebrafish reveal functional roles for the Dp in odor-evoked activity similar to those performed by the mammalian piriform cortex^60, 61^. The altered position of the Dp relative to the Dl-v is accounted for in the partial eversion model proposed by Wulliman and Mueller^2^. In support of this model, we reveal transcriptional similarity between the cichlid Dp and its associated cell-types to the mammalian piriform cortex and the corresponding region in turtles (aLC). These data are compelling evidence for molecular conservation of pallial olfactory structures across vertebrates.

In mammals, the ventral and dorsal pallial divisions give rise to the pallial amygdala and neocortex, respectively, both of which are highly complex structures whose homologues in other vertebrates have been the subject of intense debate. Several competing hypotheses have emerged, for example, some consider the sauropsid dorsal ventricular ridge (DVR) as a putative cortical homologue while others view it as amygdalar^62, 63^. Similarly, the teleost Dm has been compared to both the dorsal and ventral pallial divisions of the tetrapod brain^2, 46, 64^. The latter comparison stems in part from a demonstrated role of the Dm in emotional learning and behavior^46, 64^ and critically, these experiments largely targeted the ventromedial portion of Dm positioned along the midline^65^. Furthermore, connections of the ventromedial Dm are comparable to those of the pallial amygdala, whereas the dorsal Dm receives ascending sensory input from the preglomerular complex resembling mammalian thalamic-neocortical pathways^65^. The aforementioned partial-eversion model accounts for these observations, classifying the dorsal Dm as a dorsal pallial derivative and the ventromedial Dm as the likely ventral pallium. In line with this model, we find that cell-types concentrated in the area of ventromedial Dm showed similarity to the turtle pDVR, which previous analysis found correlated with the pallial amygdala^7^.

The origins of the mammalian six-layered neocortex and its underlying cell populations remain unclear. Structural evidence from jawless fish suggests that a precursor of the neocortex may exist in all vertebrates^66^. However, there is still no consensus as to which pallial populations in fish and sauropsids may be evolutionary precursors of neocortical populations in mammals. A variety of evidence suggests that the DVR may be functionally similar to the neocortex^62^, but an increasingly popular hypothesis is that the neocortex arose from the reptilian dorsal cortex^63^. Recent analysis of turtle scRNA-seq data found that both regions possessed cell-types with significant correlations to mammalian neocortex using all genes, but only cell-types in the anterior dorsal cortex (aDC) had significant correlations when the analysis was restricted to transcription factors^7^. In contrast, our SAMap integration revealed no significant difference in the number of transcription factors (or other gene categories) driving relationships between cell-types in the mouse neocortex and turtle aDVR/aDC. Additionally, previous analyses found that songbird cell-types most similar to the turtle aDVR (HVC_Glut-2 and HVC_Glut-5) also resembled the mammalian neocortex using both effector genes and transcription factors separately^8^. Here, we find evidence of molecular conservation between the following regions and their associated cell-types: the cichlid Dl-g, mouse neocortex, and turtle aDVR, in addition to cell-types in axolotl and songbirds which are transcriptionally similar to the turtle aDVR. In teleosts, the Dl-g is involved in visual processing^67^ and forms connections akin to those of mammalian cortical layers^49, 68^. Taken together, our results and those from previous analyses^8, 9^ suggest that neuronal cell-types transcriptionally similar to those populating turtle aDVR are conserved across vertebrates and may represent precursors to well-studied neocortical populations in the mammalian brain.

## Supporting information

Supplemental Figures and Results

Inventory of Supplement

## ACKNOWLEDGEMENTS

We dedicate this work to Karen Maruska for her groundbreaking research in teleost neuroanatomy and behavioral neuroscience. We thank the Georgia Tech Petit Institute Molecular Evolution Core for their integral roles in sequencing. This work was supported in part by NIH R01GM101095 and R01GM144560 to J.T.S., NIH F32GM128346 to Z.V.J. and Human Frontiers Science Program RGP0052/2019 to J.T.S.

## AUTHOR CONTRIBUTIONS STATEMENT

B.E.H. and G.W.G. took the lead on writing the manuscript with contributions from Z.V.J. and critical feedback from Z.V.J and J.T.S. G.W.G. took the lead on designing and creating figures with critical feedback from B.E.H., Z.V.J., and J.T.S. B.E.H. and C.M.B. optimized spatial transcriptomics wetlab protocol. Z.V.J. and B.E.H collected samples and B.E.H performed all downstream wetlab work and library preparation for spatial transcriptomics. The Petit Institute Molecular Evolution Core at GT performed sequencing for spatial transcriptomics. G.W.G. performed all bioinformatics and computational tasks, including: processing reads, quality control, clustering, marker gene analysis, prediction of the anatomical location of cell-types and cross-species gene expression comparisons. B.E.H. reviewed and synthesized literature on teleost neuroanatomy and cell-type biology and performed manual anatomical annotation of spatial transcriptomics spots. B.E.H. and G.W.G. reviewed literature on comparative vertebrate neuroanatomy. G.W.G. designed downstream statistical analyses with feedback from B.E.H., Z.V.J., and J.T.S. J.T.S. funded spatial transcriptomics experiments.

## METHODS

### Subjects

*Mchenga conophoros* cichlid species used in this study were fertilized and raised into adulthood (>180 days) in the Georgia Institute of Technology (Atlanta, GA) Engineered Biosystems Building cichlid aquaculture facilities in accordance with the Institutional Animal Care and Use Committee guidelines (IACUC protocol number A100029). All animals were collected as fry from mouthbrooding females 14 days post-fertilization and raised on a ZebTec Active Blue Stand Alone system until approximately 60 days post-fertilization, whereby animals were transferred to 190-L (92 cm long x 46 cm wide x 42 cm tall) glass aquaria ‘home tanks’ maintained on a central recirculating system. Environmental conditions of aquaria were as follows: subjects were housed in pH=8.2, 26.7°C water in social communities (20-30 mixed-sex individuals) and maintained on a 12-h:12-h light:dark cycle (full lights on between 8am-6pm Eastern Standard Time (EST)) and dim lights for 60 minute periods between light-dark transition (7am-8am and 6pm-7pm EST). Subjects were fed twice daily between 8-9am and 2-3pm (Spirulina Flake; Pentair Aquatic Ecosystems, Apopka, FL, U.S.A.). Reproductive adult subject males were introduced from home tanks to experimental tanks as described in Johnson et al. 2023^15^ which contained sand and four reproductive adult age, size-matched stimulus females of the same species.

### Tissue Processing

Adult subject males (n=2) were collected from experimental tanks between 11am-2pm EST (3-5 h after full lights-on) to control for potential effects of food intake and circadian timing on brain gene expression. Subjects were rapidly anesthetized with tricaine immediately following collection, measured for body mass (BM) and standard length (SL), and decapitated for brain extraction. Telencephala were dissected under a dissection microscope (Zeiss Stemi DV4 Stereo Microscope 8x - 32x, 000000-1018-455), in Hibernate AB Complete nutrient medium (HAB; with 2% B27 and 0.5 mM Glutamax; BrainBits) containing 0.2 U/μl RNase Inhibitor (Sigma) to prevent RNA degradation. Immediately following dissection, telencephala were embedded in disposable base cryomolds (7mm x 7mm x 5mm; Simport Scientific) containing chilled Optimal Cutting Temperature (OCT) compound (TissueTek Sakura) and flash-frozen on dry ice. Testes were then surgically extracted and weighed to calculate gonadosomatic index (GSI=gonad mass/BM*100) for each subject (subject information available in Table S1). Tissue blocks were stored in 5ml CryoELITE tissue vials (Wheaton) at −80 °C until further processing.

Telencephala were cryo-sectioned coronally at 10 μm thickness using a Cryostar NX70 cryostat at −20°C. Four tissue sections per subject were mounted onto 6.5 mm^2^ capture areas on pre-cooled Visium Spatial Gene Expression slides (10X Genomics, 200233) and slides were stored at −80°C for further processing. RNA quality of the tissue sections (RIN > 7) was confirmed using the Agilent RNA 6000 Nano Kit on the Bioanalyzer 2100 system (Agilent). Visium spatial gene expression slides were processed according to manufacturer instructions (10X Genomics; Methanol Fixation, H&E Staining, and Imaging – Visium Spatial Protocol CG000160). Slides were warmed to 37°C for 1 minute and fixed in methanol at −20°C for 30 minutes, followed by isopropanol incubation for 1 minute at room temperature. Tissue sections were then stained for hematoxylin and eosin (H&E). Brightfield images of H&E-stained slides were taken at 1.33 mm/pixel resolution using a Zeiss AxioObserver Z1 Fluorescent Microscope (Zeiss) with a 5X objective and stitched using Zen 2 software (blue edition, Zeiss) prior to library construction.

### Visium spatial gene expression library generation

Dual-index Illumina paired-end spatial gene expression libraries were prepared according to manufacturer instructions (10X Genomics; Visium Spatial Gene Expression Reagent Kits User Guide CG000239). Fixed and stained tissue sections were enzymatically permeabilized for 18 minutes. Optimal permeabilization time was determined to be 18 minutes based on initial tissue optimization trials (10X Genomics; Visium Spatial Tissue Optimization Reagent Kits User Guide CG000238). Poly-adenylated mRNA released from cells was captured by primers on the underlying spatially-barcoded 55mm-diameter gene expression spots. Primers include a 10X spatial barcode, unique molecular identifier (UMI) and an Illumina read 1 sequencing primer. Incubation with reverse transcription (RT) reagents produced spatially barcoded, full-length cDNA. Barcoded cDNA was denatured, transferred into tubes, and amplified via PCR. Amplified cDNA was size-selected with SPRIselect. During final library construction, P5 and P7 paired-end construct sequences, i5 and i7 sample indices, and an Illumina read 2 primer sequence were added. Quality was assessed using high sensitivity DNA analysis on the Bioanalyzer 2100 system (Agilent). Libraries were pooled together and sequenced on the NovaSeq 6000 (Illumina) platform using the 150-cycle SP Reagent Kit (800M reads; sample information available in Table S1).

### Spatial transcriptomics data pre-processing and quality control

FASTQ files were processed with 10x Genomics Space Ranger 1.3.1. Reads were aligned to the *Maylandia zebra* Lake Malawi cichlid genome assembly ^16^ using a splice-aware alignment algorithm (STAR) within Space Ranger and gene annotations were obtained from the same assembly (NCBI RefSeq assembly accession: GCF_000238955.4, M_zebra_UMD2a). H&E images were oriented so fiducial markers were in the correct location and the appropriate slide layout files were chosen. Barcoded spot patterns were aligned to the input slide image and tissue was distinguished from background on the slide. Following these steps, Space Ranger generated filtered feature-barcode matrices (one per slide) containing expression data for a total of 32,471 features (corresponding to annotated genes) and a total of 6,755 barcodes (corresponding to spots). In the R the “Seurat” package was used to remove spots with less than one UMI and spots containing small pieces of stray tissue (48 spots removed).

### Selection of tissue sections for downstream analysis

This analysis focuses on seven tissue sections from two subject males, S1 (n = 3 sections) and S2 (n = 4 sections). This decision was made based on 1) the high quality of the tissue sections, and 2) the observation that the combination of S1 and S2 sections yield a comprehensive representation of telencephalic subregions along the rostral-caudal axis. The order of the tissue hemispheres along the rostral-caudal axis was determined through visual inspection of H&E-stained tissue images.

### Clustering and selection of parameters

To prevent over-tuning of clustering parameters, a systemic method was used to determine optimal clustering parameters. ChooseR is a tool compatible with Seurat that evaluates clustering quality based on robustness metrics of bootstraps of the data. The code was adapted to evaluate more clustering parameters and to follow a slightly different workflow. The additional parameters that were evaluated were min.dist and n.neighbors in the RunUMAP function from Seurat. To ensure that bootstraps were entirely independent, the find_clusters function was modified to include the SCTransform, RunPCA, and RunUMAP functions from Seurat. Without these functions, there would be information leakage in the form of the variable genes and reduced dimensional space. In SCTransform, tissue sections were regressed out using var.to.regress and the resulting “SCT” assay was set as the active assay. 50 dimensions were used in RunPCA and RunUMAP. Additionally, in RunUMAP spread was set to 1, n.epochs was set to 1000 and metric was set to “euclidean”. In FindNeighbors, reduction was set to “UMAP” and the first two dimensions were used, k.param was set to the same value as n.neighbors, n.trees was set to 500, and prune.SNN was set to 0. In Find Clusters, algorithm was set to 2.

ChooseR was used to evaluate the clustering of combinations of min.dist and n.neighbors in RunUMAP and resolution in FindClusters. The set of values tested were as follows: min.dist of 0.1, 0.2, 0.3, 0.4 and 0.5; n.neighbors of 10, 20, 30, 40 and 50; and resolution of 0.2, 0.4, 0.6, 0.8, 1.0 and 1.2. The default value of 80% subsampling of the data was used as well as the default of 100 bootstraps (per combination of parameters). ChooseR defines the near-optimal clustering parameters as the one yielding the highest number of clusters whose median silhouette score greater than with the highest lower bound (95% confidence interval). Using this same methodology for our set of clustering parameters, resulted in min.dist=0.1, n.neighbors=30, and resolution=1.2.

### Human Ortholog identification

Teleosts including *M.zebra* have undergone a genome duplication since their divergence with mammals, thus identification of human orthologs is non-trivial and multiple methods were used to ensure accuracy. Human orthologs to cichlid genes were retrieved from ENSEMBL and Varadarajan et al 2018^69^, which uses a protein based approach. To perform interspecies correlation analysis, one-to-one orthologs are required. For all orthologs, cichlid genes were kept only where the human ortholog was contained in one of the following: NCBI gene name, NCBI gene symbol, ENSEMBL gene symbol, Varadarajan et al 2018^69^, and human ortholog from ENSEMBL. Due to the aforementioned genome duplication, there are numerous many-to-one orthologs and discarding them all would leave few genes remaining. Thus, in cases of many-to-one orthologs the cichlid gene with the greatest consistency among the aforementioned sources was kept and contained the greatest number of UMIs in the spatial transcriptomics and single nuclei gene expression matrices, while the others were removed. This resulted in 13,237 cichlid genes. For interspecies integration analysis, a different method, BLAST, was used for ortholog identification (see ‘Interspecies integration of single cell/nucleus RNA-seq data’).

### Computation of cell-type abundance estimation in spatial transcriptomics spots

For cell-type deconvolution of spots, cell2location^32^ was used as it was among the top performing tools for this task reported by a recent benchmarking paper^33^. Cell2location is a Bayesian model that estimates the absolute abundance of cell-types at each spot. First, the regression model for the single cell data was initialized with default settings, using batch as the batch_key and the model was trained using a maximum of 250 epochs. Next, the regression model for the spatial transcriptomics data was initialized with the single cell reference signatures, default settings, and hyperparameters selected based on cell2location’s recommendations. We did not observe strong within-batch variation in total RNA count, therefore we set detection_alpha=200. By manual visual inspection, we estimated the number of cells per spot to be 8 and set N_cells_per_location=8. The model was then trained using a maximum of 30,000 epochs. Finally, the estimated mean abundance of cell-types was rounded and spots with no estimated cells were assigned one cell to the cell-type with greatest abundance.

### Manual cell abundance estimation in spatial transcriptomics spots

To evaluate the accuracy of the estimates of cell abundance in spatial spots by cell2location, cell bodies were manually counted for a select number of spots in a single tissue hemisphere (S_1_C_2_R). Due to a high density of cell bodies at the peripheral edges of the tissue where individual cell bodies could not be reliably distinguished, these spots were not manually counted and excluded from the comparison. The Pearson’s and Spearman’s correlation coefficient was found between manual and computational estimates of cell abundance (Fig. S7). Note that cell bodies stained and visible for manual estimation may not reflect the same biological component as was computationally estimated.

### scRNA-seq dataset retrieval

Mouse gene expression data was retrieved from Zeisel et al 2018^36^ and Saunders et al 2018^44^. While gene expression data for other vertebrate taxa was retrieved from Lust et al 2022^9^ (axolotl), Tosches et al 2018^7^ (turtle) and Colquitt et al 2022^8^ (songbirds). Next, the datasets were filtered to include only cells from the telencephalon. Data from Zeisel et al was filtered to include only cells from cell-types annotated as belonging to the telencephalon and data from Saunders et al was only retrieved for the following brain regions: Frontal Cortex, Globus Pallidus, Hippocampus, Posterior Cortex, and Striatum. No filtering was required for the axolotl and turtle datasets, as the cells came from the telencephalon. All cells from the HVC and RA of songbirds were kept, but only MSNs from Area X were retained due to non-descriptive annotations of the other cells. Next, cells annotated by Zeisel et al as low-quality were removed. This information was not readily available for Saunders et al, thus the dataset was subset by cells with greater than 500 UMIs, 500 genes, less than 5000 UMIs and less than 2500 genes. The other vertebrate datasets retrieved were already filtered, so no further filtering was performed. Finally, due to the large number of cells from Saunders et al, cells were randomly and evenly removed from the largest clusters until 100,000 cells remained. All datasets were normalized using SCTransform using default options.

### Interspecies correlation of single cell/nucleus RNA-seq data

To compare sc/snRNA-seq from cichlid and mouse telencephalon, a procedure similar to Tosches et al 2018^7^, Colquitt et al 2022^8^ and Lust et al 2022^9^ was implemented as it is well suited for comparisons of distantly related species. Briefly, a Spearman correlation and p-value are found from the average expression of DEGs present in both species. For robustness, correlations were calculated using multiple mouse datasets^36, 44^. Interspecies correlations from Zeisel et al 2018^36^ were performed on clusters. Likewise correlations were calculated on clusters and subclusters with data from Saunders et al^44^.

DEGs were found in the cichlid and mouse datasets using the FindAllMarkers functions with only.pos = T and subsetting by results with adjusted p-values less than 0.05. Then, the common set of genes that were DEGs in both species were found (see ‘Human Ortholog identification’) and their average gene expression was calculated across the whole dataset using the AverageExpression function in Seurat. Similar to Colquitt et al 2022, these average gene expression values were normalized by log(x+1) + 0.1 and were divided by the mean for each gene across the dataset. The normalized average gene expression matrices for the cichlid and mouse datasets were then correlated using a Spearman correlation. Significance of these correlations was determined using two criteria: 1) Bonferroni adjusted p-value of Spearman correlation test < 0.05 and 2) Bonferonni adjusted p-value of permutation test < 0.05. The former was calculated using cor.test in R with method = “spearman” and alternative = “greater”. Similar to Tosches et al 2018^7^, the latter was found by shuffling the normalized average gene expression values across cell-types in the cichlid gene by cluster matrix, then re-calculating the Spearman correlations with the mouse gene by cluster matrix. 1,000 permutations were performed and the p-value was calculated by finding the proportion of correlations from the permutations greater than the real, non-permuted correlations.

### Interspecies integration of single cell/nucleus RNA-seq data

Integration of sc/snRNA-seq data from vertebrate taxa was performed to compare the transcriptional similarities of their cell-types. To achieve this, SAMap^13^, a python package, was chosen as it was designed for comparative analyses of distantly related species. SAMap considers sequence similarity of genes between species in its projection of the datasets in joint lower-dimensional spaces. Allowing genes with similar sequences to weigh more heavily in the integration, which is advantageous when comparing species with large evolutionary distances like cichlids and mice. First, to determine sequence similarity, the proteomes of the vertebrate taxa were downloaded from NCBI using the version of the assemblies used to create the sc/snRNA-seq datasets: *Maylandia zebra* (GCF_000238955.4), *Mus musculus* (GCF_000001635.27), *Chrysemys picta bellii* (GCF_000241765.3) and *Taeniopygia guttata* (GCF_003957565.1). For the axolotl (*Ambystoma mexicanum*) dataset, the assembly was retrieved from v6.0 release of the genome not located on NCBI (AmexT_v47). Next, blastp was executed to identify reciprocal blast hits between vertebrate taxa and cichlids using SAMap’s map_genes.sh script. Then, the raw gene by counts matrix of the sc/snRNA datasets from their Seurat objects were saved as an h5ad file and processed by the SAMAP function with arguments specifying the gene symbols of each protein and the directory containing the blastp results. Finally, the run function was then executed using default parameters.

The knn-graph produced by SAMap was then used to create a similarity score between cell-types in cichlids and other vertebrates. The similarity score was defined as the mean number of k-nearest-neighbors from other vertebrates to cichlid nuclei and to determine significance, a permutation test was performed. The cell-type labels of cells/nuclei for both datasets were shuffled and similarity scores were calculated. This was repeated 1,000 times and cross-species cell-type pairs with similarity scores greater than all permutations of each cichlid cell-type were considered significant.

### Identification of genes driving cell-type relationships discovered using SAMap

To find the genes driving cell-type relationships, the function GenePairFinder from SAMap was used with default parameters. Then for every combination of cell-types, the function find_genes function from SAMap was used to find the genes driving the relationship, with n_genes set to the total number of genes in the datasets. The default parameter returns the top 1,000 genes driving the relationship, but we allowed the function to return all the genes it found to drive the relationship. These genes are defined by SAMap as genes that contribute positively to the correlation of the cell-types and are differentially expressed in each cell-type.

### Identification of genes upregulated in conserved cell-types across vertebrates

Marker genes (significantly differentially expressed genes with p_val_adj <0.05, log_2_FC > 0 and min.pct > 0.1) with human orthologs present in all datasets analysis for select strongly conserved cell-types were found. Strongly conserved cell-types included: OLIG (mouse OEC; bird Oligo; turtle tsOlig; axolotl OLIG15; cichlid 2.2_Oligo), OB (mouse OBDOP2; bird GABA-1-1; turtle i01; axolotl GABA3; cichlid 5.2_GABA), MGE1 (mouse TEINH17; bird GABA-3; turtle i07; axolotl GABA2; cichlid 6_GABA), MGE2 (mouse TEINH21; bird GABA-2; turtle i08; axolotl GABA17; cichlid 15.3_GABA), MSN (mouse MSN1; bird MSN3; turtle i05; axolotl GABA11; cichlid 4.1_GABA), DGNBL (mouse DGNBL1; bird Pre-2; turtle tsNPCs; axolotl NB1; cichlid 9.5_Glut) and CA3 (mouse TEGLU23; bird HVC_Glut-3; turtle e34; axolotl GLUT7; cichlid 8.9_Glut). Additionally, human orthologs in conserved cell-types that were upregulated (positive log_2_FC value), but not meeting the other criteria of marker genes were found. When multiple genes in a species had the same human ortholog, the gene with the greatest log_2_FC was selected.

### Interspecies comparison of anatomical regions

To gain insight into the similarity of anatomical regions across vertebrates, we compared scRNA-seq data from turtles and spatial transcriptomics data from mice to cichlid spatial transcriptomics data separately. Both analyses were performed in the same manner described in ‘Interspecies integration of single cell/nucleus RNA-seq data’. For the comparison to the turtle dataset, the detailed anatomical labels were used when available and the broad anatomical labels were used when it was not. The mouse spatial transcriptomics dataset was retrieved from Ortiz et al 2020^45^, the dataset was subset to include only cells from the cerebrum (Allen Brain Atlas abbreviation ‘CH’), the ventricular systems (Allen Brain Atlas abbreviation ‘VS’), and fiber tracts (Allen Brain Atlas abbreviation ‘fiber tracts’). Then, the dataset was normalized using SCTransform, and the annotations from ABA were used (ABA_acronym column in the metadata). Since these annotations were extremely detailed, the parent annotation was used in some cases. Moreover, spots annotated as cortex (“CTX”) with no more detail were assigned to “U_CTX” to represent undefined cortical regions.

### Composition of genes driving effects and conserved marker genes by transcription factors, neuromodulatory ligands and receptors

To determine differences in the shared marker genes between cell-types with significant similarities across species, the composition of the genes driving effects and shared markers by transcription factors, neuromodulatory ligands and receptors was found. Transcription factors were obtained from Tosches et al 2018^7^ and neuromodulatory ligands and receptors were obtained from Johnson et al 2023^15^. Since the aforementioned lists use human orthologs, the human ortholog of genes driving effects were found and the human ortholog of mouse genes were used when possible (ie in cichlid-mouse and turtle-mouse comparisons) and otherwise the human ortholog of cichlid genes were used (ie cichlid-turtle comparison). The percentage of genes in these lists was calculated and compared for across cell-type hits. To determine significance, a χ^2^ test was performed on the mean percentage of transcription factors, neuromodulatory ligands and receptors.

## Notes

### Competing Interest Statement

The authors have declared no competing interest.

## REFERENCES

1. O’Connell, L. A. & Hofmann, H. A. The vertebrate mesolimbic reward system and social behavior network: a comparative synthesis. J. Comp. Neurol. 519, 3599–3639 (2011).

2. Wullimann, M. F. & Mueller, T. Teleostean and mammalian forebrains contrasted: Evidence from genes to behavior. J. Comp. Neurol. 475, 143–162 (2004).

3. Yamamoto, N. et al. A New Interpretation on the Homology of the Teleostean Telencephalon Based on Hodology and a New Eversion Model. Brain, Behavior and Evolution vol. 69 96–104 Preprint at https://doi.org/10.1159/000095198 (2007).

4. Braford, M. R., Jr. Comparative aspects of forebrain organization in the ray-finned fishes: touchstones or not? Brain Behav. Evol. 46, 259–274 (1995).

5. Northcutt, R. G. Connections of the lateral and medial divisions of the goldfish telencephalic pallium. J. Comp. Neurol. 494, 903–943 (2006).

6. Nieuwenhuys, R. The development and general morphology of the telencephalon of actinopterygian fishes: synopsis, documentation and commentary. Brain Struct. Funct. 215, 141–157 (2011).

7. Tosches, M. A. et al. Evolution of pallium, hippocampus, and cortical cell types revealed by single-cell transcriptomics in reptiles. Science 360, 881–888 (2018).

8. Colquitt, B. M., Merullo, D. P., Konopka, G., Roberts, T. F. & Brainard, M. S. Cellular transcriptomics reveals evolutionary identities of songbird vocal circuits. Science 371, (2021).

9. Lust, K. et al. Single-cell analyses of axolotl telencephalon organization, neurogenesis, and regeneration. Science 377, eabp9262 (2022).

10. Woych, J. et al. Cell-type profiling in salamanders identifies innovations in vertebrate forebrain evolution. Science 377, eabp9186 (2022).

11. Hain, D. et al. Molecular diversity and evolution of neuron types in the amniote brain. Science 377, eabp8202 (2022).

12. Pandey, S., Moyer, A. J. & Thyme, S. B. A single-cell transcriptome atlas of the maturing zebrafish telencephalon. Genome Res. 33, 658–671 (2023).

13. Tarashansky, A. J. et al. Mapping single-cell atlases throughout Metazoa unravels cell type evolution. (2021) doi:10.7554/eLife.66747.

14. Styfhals, R. et al. Cell type diversity in a developing octopus brain. Nat. Commun. 13, 7392 (2022).

15. Johnson, Z. V. et al. Cellular profiling of a recently-evolved social behavior. bioRxiv 2022.08.09.503380 (2022) doi:10.1101/2022.08.09.503380.

16. Conte, M. A. et al. Chromosome-scale assemblies reveal the structural evolution of African cichlid genomes. Gigascience 8, (2019).

17. Ganz, J. et al. Subdivisions of the adult zebrafish subpallium by molecular marker analysis. J. Comp. Neurol. 520, 633–655 (2012).

18. Ganz, J. et al. Subdivisions of the adult zebrafish pallium based on molecular marker analysis. F1000Research vol. 3 308 Preprint at https://doi.org/10.12688/f1000research.5595.2 (2015).

19. Maruska, K. P., Butler, J. M., Field, K. E. & Porter, D. T. Localization of glutamatergic, GABAergic, and cholinergic neurons in the brain of the African cichlid fish, Astatotilapia burtoni. J. Comp. Neurol. 525, 610–638 (2017).

20. Diotel, N. et al. Comprehensive expression map of transcription regulators in the adult zebrafish telencephalon reveals distinct neurogenic niches. J. Comp. Neurol. 523, 1202–1221 (2015).

21. Distributions and colocalization of neuropeptide Y and somatostatin in the goldfish brain. J. Chem. Neuroanat. 5, 221–233 (1992).

22. Grone, B. P., Butler, J. M., Wayne, C. R. & Maruska, K. P. Expression patterns and evolution of urocortin and corticotropin-releasing hormone genes in a cichlid fish. Journal of Comparative Neurology vol. 529 2596–2619 Preprint at https://doi.org/10.1002/cne.25113 (2021).

23. Yamamoto, K., Ruuskanen, J. O., Wullimann, M. F. & Vernier, P. Differential expression of dopaminergic cell markers in the adult zebrafish forebrain. J. Comp. Neurol. 519, 576–598 (2011).

24. O’Connell, L. A., Fontenot, M. R. & Hofmann, H. A. Characterization of the dopaminergic system in the brain of an African cichlid fish, Astatotilapia burtoni. The Journal of Comparative Neurology vol. 519 75–92 Preprint at https://doi.org/10.1002/cne.22506 (2011).

25. Adolf, B. et al. Conserved and acquired features of adult neurogenesis in the zebrafish telencephalon. Dev. Biol. 295, 278–293 (2006).

26. Gokce, O. et al. Cellular Taxonomy of the Mouse Striatum as Revealed by Single-Cell RNA-Seq. Cell Rep. 16, 1126–1137 (2016).

27. Aoki, T. et al. Imaging of neural ensemble for the retrieval of a learned behavioral program. Neuron 78, 881–894 (2013).

28. Rink, E. & Wullimann, M. F. The teleostean (zebrafish) dopaminergic system ascending to the subpallium (striatum) is located in the basal diencephalon (posterior tuberculum). Brain Research vol. 889 316–330 Preprint at https://doi.org/10.1016/s0006-8993(00)03174-7 (2001).

29. Northcutt, R. G. Forebrain evolution in bony fishes. Brain Res. Bull. 75, 191–205 (2008).

30. Mueller, T., Dong, Z., Berberoglu, M. A. & Guo, S. The dorsal pallium in zebrafish, Danio rerio (Cyprinidae, Teleostei). Brain Res. 1381, 95–105 (2011).

31. Alderman, S. L. & Bernier, N. J. Localization of corticotropin-releasing factor, urotensin I, and CRF-binding protein gene expression in the brain of the zebrafish, Danio rerio. J. Comp. Neurol. 502, 783–793 (2007).

32. Kleshchevnikov, V. et al. Cell2location maps fine-grained cell types in spatial transcriptomics. Nat. Biotechnol. 40, 661–671 (2022).

33. Li, B. et al. Benchmarking spatial and single-cell transcriptomics integration methods for transcript distribution prediction and cell type deconvolution. Nat. Methods 19, 662–670 (2022).

34. York, R. A., Patil, C., Darrin Hulsey, C., Todd Streelman, J. & Fernald, R. D. Evolution of bower building in Lake Malawi cichlid fish: phylogeny, morphology, and behavior. Frontiers in Ecology and Evolution vol. 3 Preprint at https://doi.org/10.3389/fevo.2015.00018 (2015).

35. Volff, J.-N. Genome evolution and biodiversity in teleost fish. Heredity 94, 280–294 (2004).

36. Zeisel, A. et al. Molecular Architecture of the Mouse Nervous System. Cell 174, 999–1014.e22 (2018).

37. Ghaddar, B., Lübke, L., Couret, D., Rastegar, S. & Diotel, N. Cellular Mechanisms Participating in Brain Repair of Adult Zebrafish and Mammals after Injury. Cells 10, (2021).

38. Grupp, L., Wolburg, H. & Mack, A. F. Astroglial structures in the zebrafish brain. J. Comp. Neurol. 518, 4277–4287 (2010).

39. März, M. et al. Heterogeneity in progenitor cell subtypes in the ventricular zone of the zebrafish adult telencephalon. Glia 58, 870–888 (2010).

40. Ming, G.-L. & Song, H. Adult neurogenesis in the mammalian central nervous system. Annu. Rev. Neurosci. 28, 223–250 (2005).

41. Hodge, R. D. et al. Tbr2 expression in Cajal-Retzius cells and intermediate neuronal progenitors is required for morphogenesis of the dentate gyrus. J. Neurosci. 33, 4165–4180 (2013).

42. Mueller, T., Wullimann, M. F. & Guo, S. Early teleostean basal ganglia development visualized by zebrafish Dlx2a, Lhx6, Lhx7, Tbr2 (eomesa), and GAD67 gene expression. J. Comp. Neurol. 507, 1245–1257 (2008).

43. Butler, A. B. & Hodos, W. Comparative Vertebrate Neuroanatomy: Evolution and Adaptation. (John Wiley & Sons, 2005).

44. Saunders, A. et al. Molecular Diversity and Specializations among the Cells of the Adult Mouse Brain. Cell 174, 1015–1030.e16 (2018).

45. Ortiz, C. et al. Molecular atlas of the adult mouse brain. Sci Adv 6, eabb3446 (2020).

46. Portavella, M., Vargas, J. P., Torres, B. & Salas, C. The effects of telencephalic pallial lesions on spatial, temporal, and emotional learning in goldfish. Brain Res. Bull. 57, 397–399 (2002).

47. Broglio, C. et al. Hippocampal Pallium and Map-Like Memories through Vertebrate Evolution. Journal of Behavioral and Brain Science vol. 05 109–120 Preprint at https://doi.org/10.4236/jbbs.2015.53011 (2015).

48. Broglio, C., Rodríguez, F., Gómez, A., Arias, J. L. & Salas, C. Selective involvement of the goldfish lateral pallium in spatial memory. Behav. Brain Res. 210, 191–201 (2010).

49. Yamamoto, N. & Ito, H. Visual, lateral line, and auditory ascending pathways to the dorsal telencephalic area through the rostrolateral region of the lateral preglomerular nucleus in cyprinids. The Journal of Comparative Neurology vol. 508 615–647 Preprint at https://doi.org/10.1002/cne.21717 (2008).

50. Murakami, T., Morita, Y. & Ito, H. Extrinsic and intrinsic fiber connections of the telencephalon in a teleost,Sebastiscus marmoratus. The Journal of Comparative Neurology vol. 216 115–131 Preprint at https://doi.org/10.1002/cne.902160202 (1983).

51. Kyle, A. L. & Peter, R. E. Effects of forebrain lesions on spawning behaviour in the male goldfish. Physiol. Behav. 28, 1103–1109 (1982).

52. Kondo, Y., Shinoda, A., Yamanouchi, K. & Arai, Y. Role of septum and preoptic area in regulating masculine and feminine sexual behavior in male rats. Horm. Behav. 24, 421–434 (1990).

53. Goodson, J. L. & Adkins-Regan, E. Effect of intraseptal vasotocin and vasoactive intestinal polypeptide infusions on courtship song and aggression in the male zebra finch (Taeniopygia guttata). J. Neuroendocrinol. 11, 19–25 (1999).

54. Font, C., Lanuza, E., Martinez-Marcos, A., Hoogland, P. V. & Martinez-Garcia, F. Septal complex of the telencephalon of lizards: III. Efferent connections and general discussion. J. Comp. Neurol. 401, 525–548 (1998).

55. Striedter, G. F. Evolution of the hippocampus in reptiles and birds. J. Comp. Neurol. 524, 496–517 (2016).

56. Bienkowski, M. S. et al. Integration of gene expression and brain-wide connectivity reveals the multiscale organization of mouse hippocampal networks. Nat. Neurosci. 21, 1628–1643 (2018).

57. van Strien, N. M., Cappaert, N. L. M. & Witter, M. P. The anatomy of memory: an interactive overview of the parahippocampal-hippocampal network. Nat. Rev. Neurosci. 10, 272–282 (2009).

58. Rodríguez, F., et al. Conservation of Spatial Memory Function in the Pallial Forebrain of Reptiles and Ray-Finned Fishes. The Journal of Neuroscience vol. 22 2894–2903 Preprint at https://doi.org/10.1523/jneurosci.22-07-02894.2002 (2002).

59. Durán, E., Ocaña, F. M., Broglio, C., Rodríguez, F. & Salas, C. Lateral but not medial telencephalic pallium ablation impairs the use of goldfish spatial allocentric strategies in a ‘hole-board’ task. Behavioural Brain Research vol. 214 480–487 Preprint at https://doi.org/10.1016/j.bbr.2010.06.010 (2010).

60. Yaksi, E., von Saint Paul, F., Niessing, J., Bundschuh, S. T. & Friedrich, R. W. Transformation of odor representations in target areas of the olfactory bulb. Nat. Neurosci. 12, 474–482 (2009).

61. Blumhagen, F. et al. Neuronal filtering of multiplexed odour representations. Nature 479, 493–498 (2011).

62. Striedter, G. F. The telencephalon of tetrapods in evolution. Brain Behav. Evol. 49, 179– 213 (1997).

63. Aboitiz, F., Morales, D. & Montiel, J. The evolutionary origin of the mammalian isocortex: towards an integrated developmental and functional approach. Behav. Brain Sci. 26, 535–52; discussion 552–85 (2003).

64. Nieuwenhuys, R. The forebrain of actinopterygians revisited. Brain Behav. Evol. 73, 229– 252 (2009).

65. Yamamoto, N. & Ito, H. Fiber connections of the anterior preglomerular nucleus in cyprinids with notes on telencephalic connections of the preglomerular complex. J. Comp. Neurol. 491, 212–233 (2005).

66. Suryanarayana, S. M., Robertson, B., Wallén, P. & Grillner, S. The Lamprey Pallium Provides a Blueprint of the Mammalian Layered Cortex. Curr. Biol. 27, 3264–3277 (2017).

67. Yamane, Y., Yoshimoto, M. & Ito, H. Area dorsalis pars lateralis of the telencephalon in a teleost (Sebastiscus marmoratus) can be divided into dorsal and ventral regions. Brain Behav. Evol. 48, 338–349 (1996).

68. Ito, H. & Yamamoto, N. Non-laminar cerebral cortex in teleost fishes? Biol. Lett. 5, 117–121 (2009).

69. Ramakrishnan Varadarajan, A., Mopuri, R., Streelman, J. T. & McGrath, P. T. Genome-wide protein phylogenies for four African cichlid species. BMC Evol. Biol. 18, 1 (2018).

