## Supplemental Figures and Results for "Spatially resolved cell atlas of the teleost telencephalon and deep homology of the vertebrate forebrain"

**SUPPLEMENT**

FIGURES

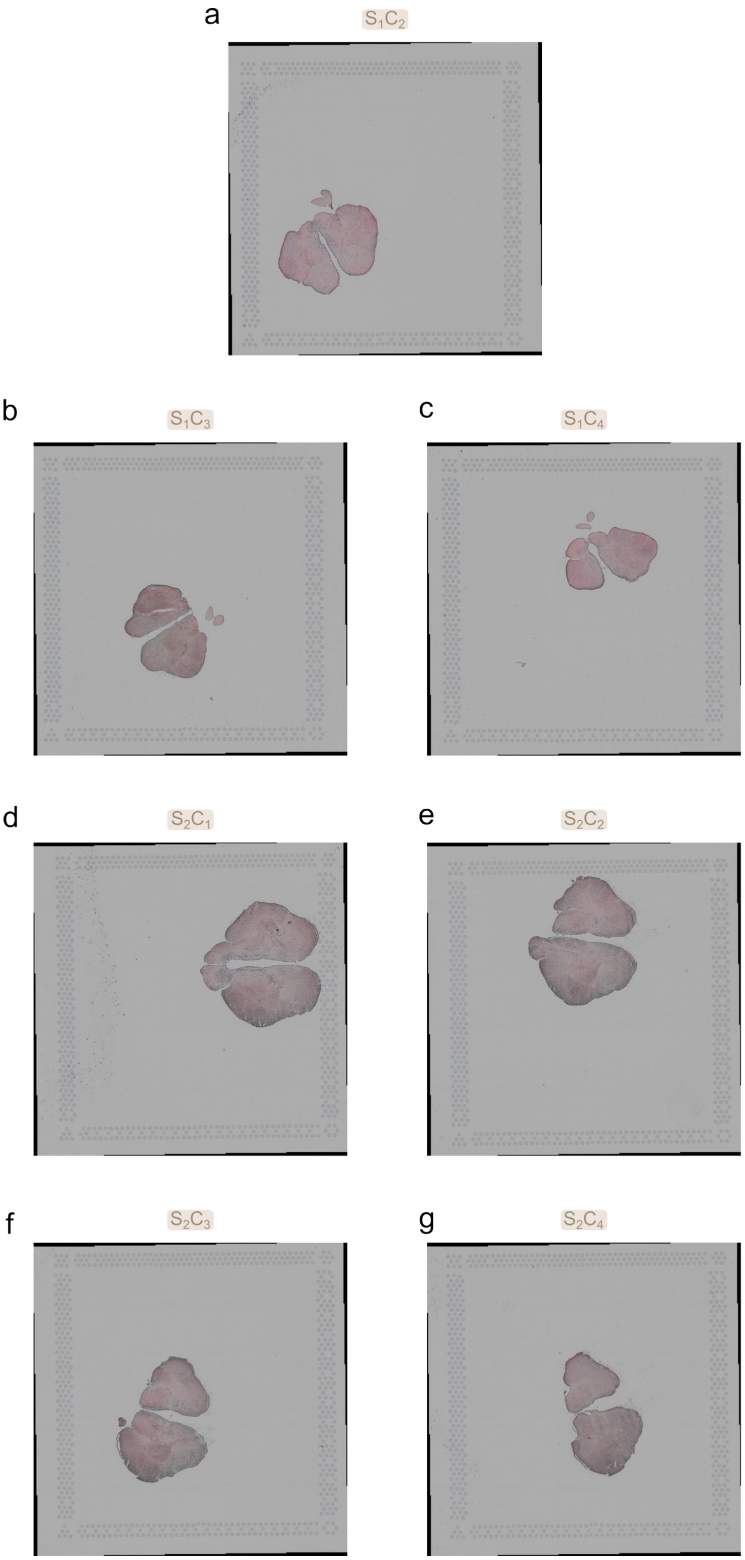


**Figure S1.** A-G. Brightfield images of H&E-stained slides for each capture area from each subject (S1C2-4 and S2C1-4), respectively.

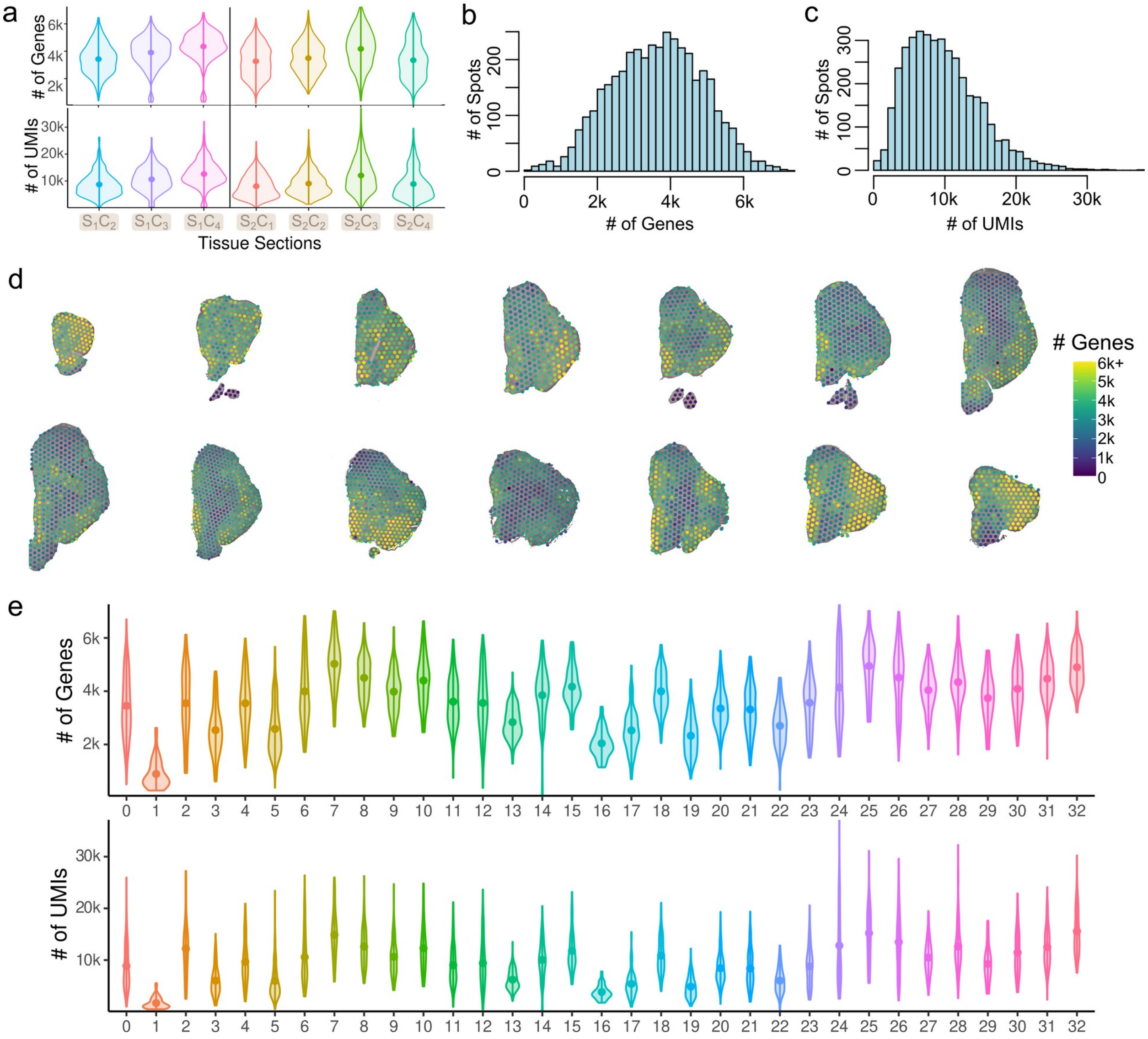


**Figure S2. Spatial transcriptomics data quality.** A. Distribution of the number of genes and UMIs across capture areas from subjects represented by a violin plot. Dots signify the mean value per capture area and vertical lines represent the range. B-C. Distribution of number of genes and UMIs across all subjects, respectively. D. The number of genes expressed in each spot shown on the tissue. E. Distribution of the number of genes and UMIs across clusters represented by a violin plot. Dots signify the mean value per cluster and vertical lines represent the range.

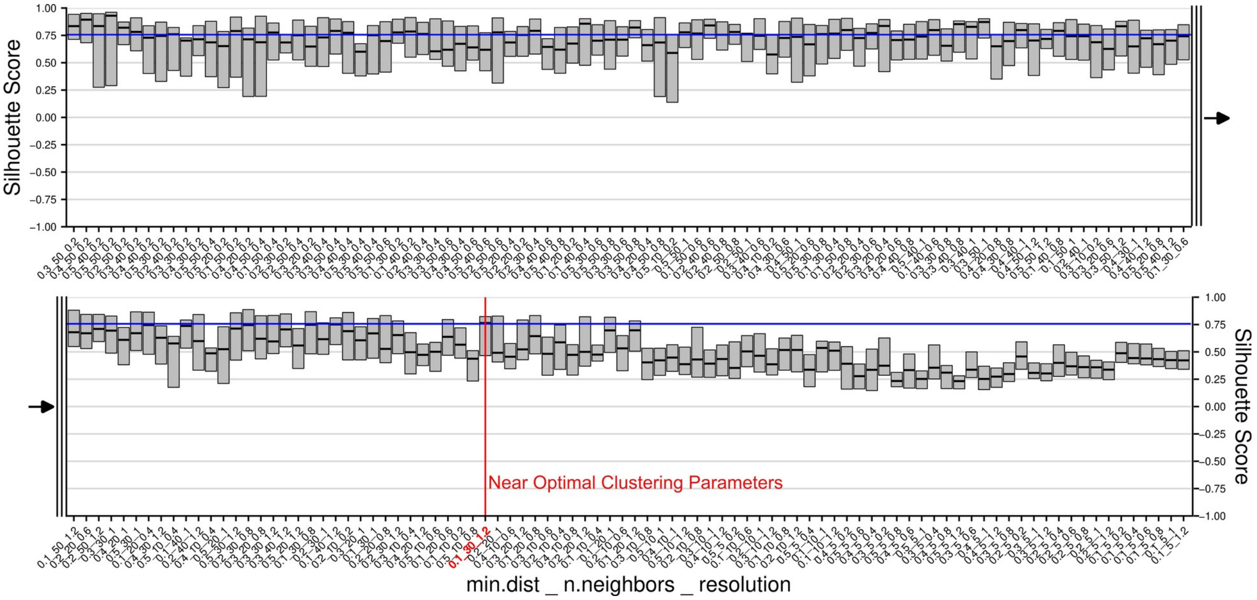


**Figure S3. Near-optimal clustering parameters of spatial transcriptomics data.** Silhouette scores shown on the y-axis for combinations of clustering parameters shown on the x-axis. Boxplots represent the silhouette scores of bootstraps of the data with medians shown as lines and shaded regions as the 95% confidence interval. ChooseR determines the near-optimal combination (red line) as the one yielding the highest number of clusters whose median silhouette score is greater than the highest lower bound of the 95% confidence interval (blue line). Combinations of values were evaluated for min.dist, n.neighbors and resolution (see Methods).

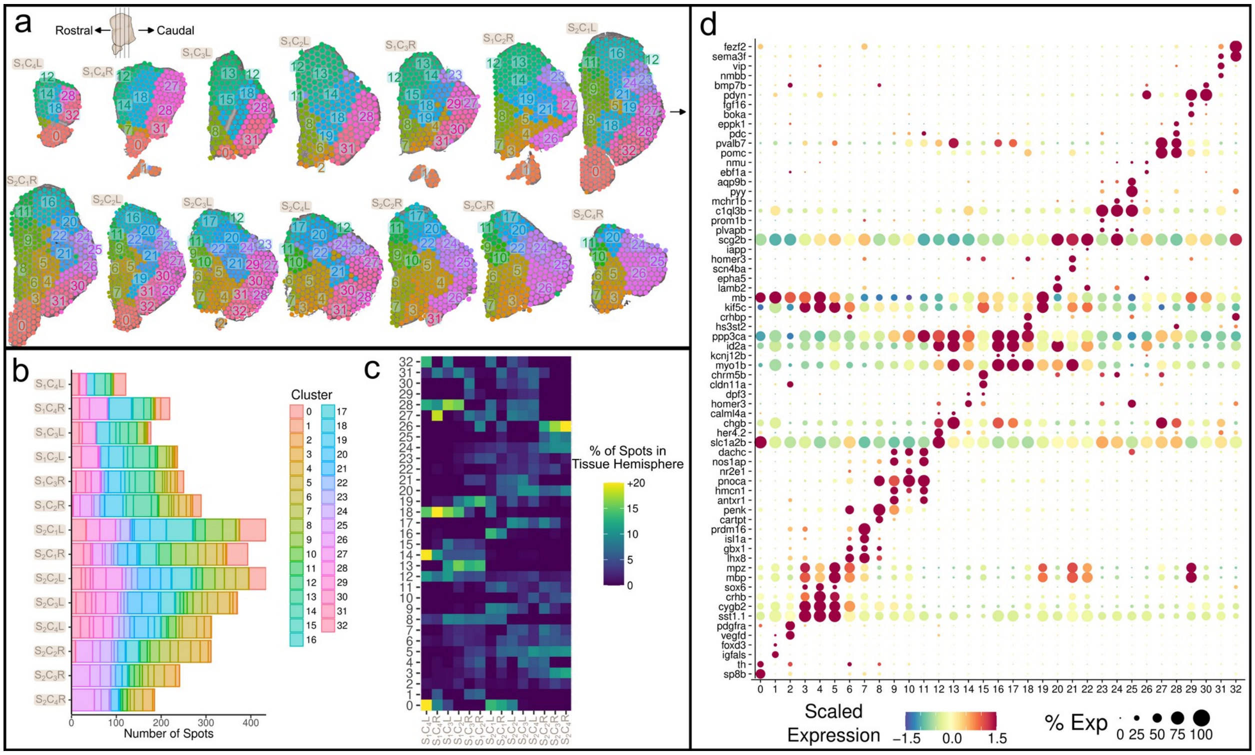

**Figure S4. Unbiased clusters of spatial transcriptomics data.** A. Spots on tissue colored by clusters with tissue hemispheres ordered rostrocaudally. B. Number of spots from each cluster in tissue hemispheres. C. Composition of spots from tissue hemispheres by clusters. D. Select top marker genes of clusters. Dots are colored by their mean scaled expression in each cluster and dot size indicates the percent of cells in clusters expressing the marker.

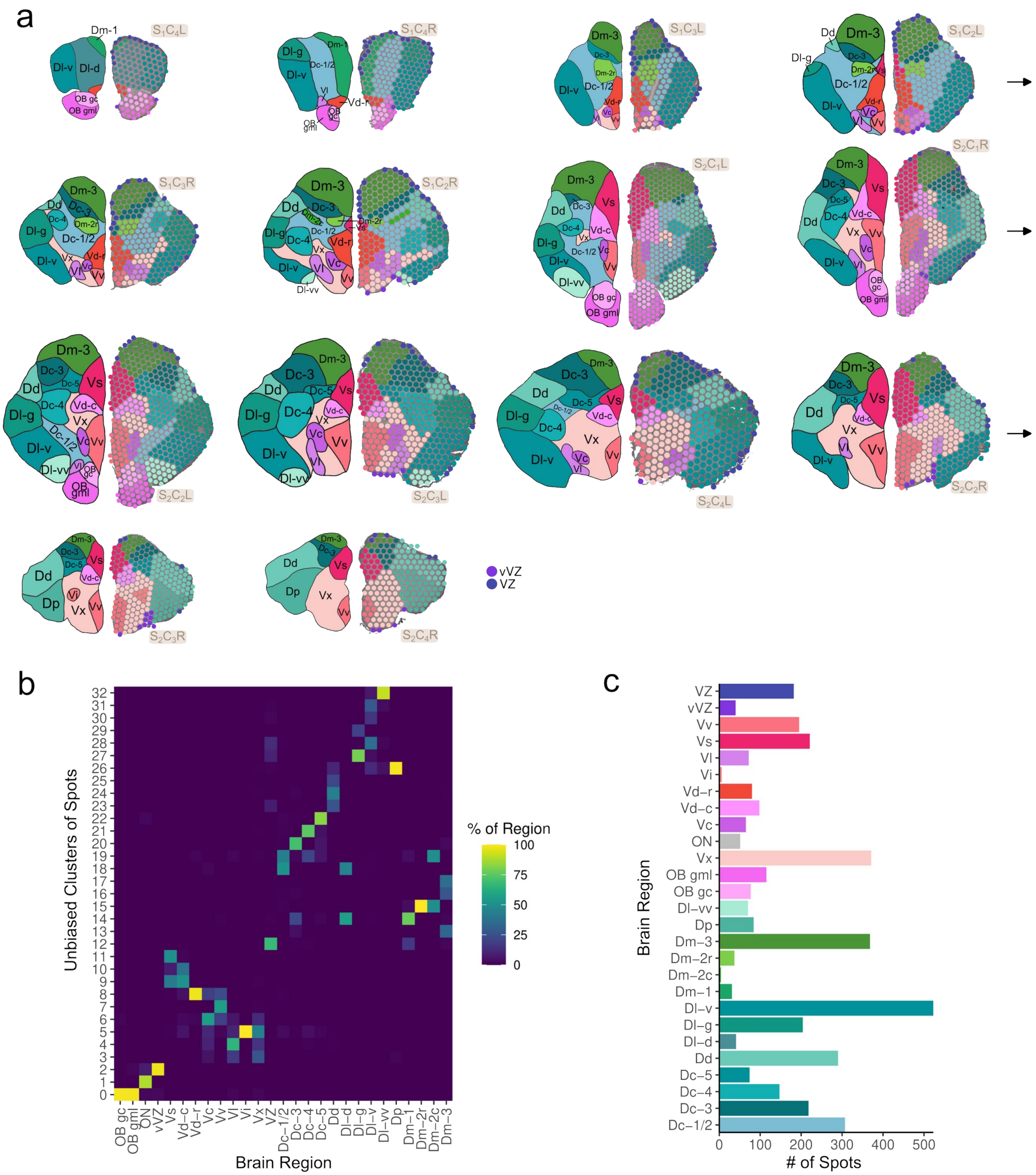

**Figure S5. Manually annotated anatomical labels of spots.** A. Annotations of anatomical subregions of the teleost telencephalon on tissue images. Each tissue hemisphere is represented on the left by an illustration of the tissue with anatomical subregions labeled and on the right by the tissue images with spots colored by anatomical labels. B. Composition of anatomical subregions by unbiased clusters. C. Number of spots per anatomical subregions.

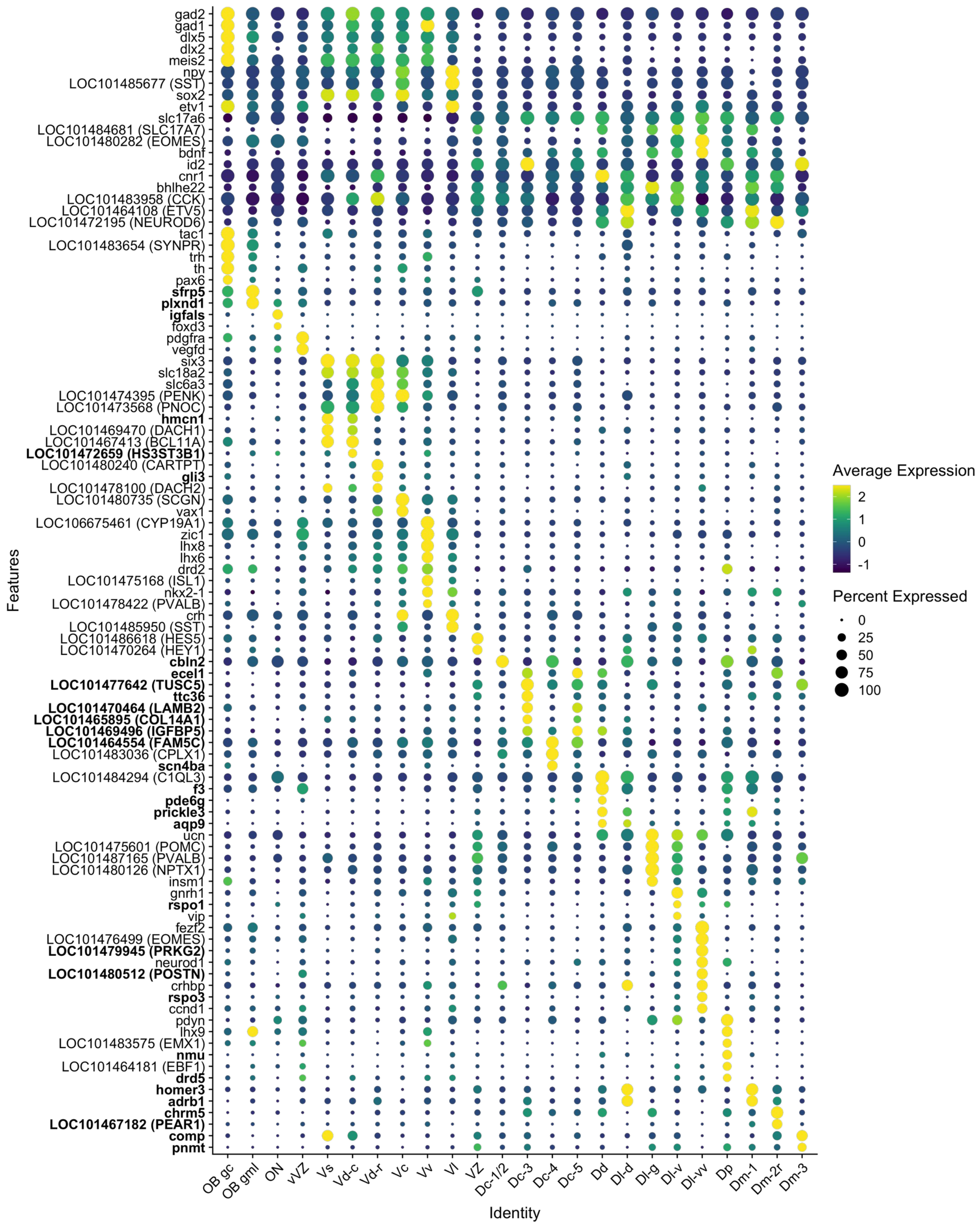


**Figure S6. Gene expression of marker genes across anatomical regions in the cichlid telencephalon.** Dots are colored by the average gene expression in each anatomical region and size indicates the percent of spots expressing the gene.

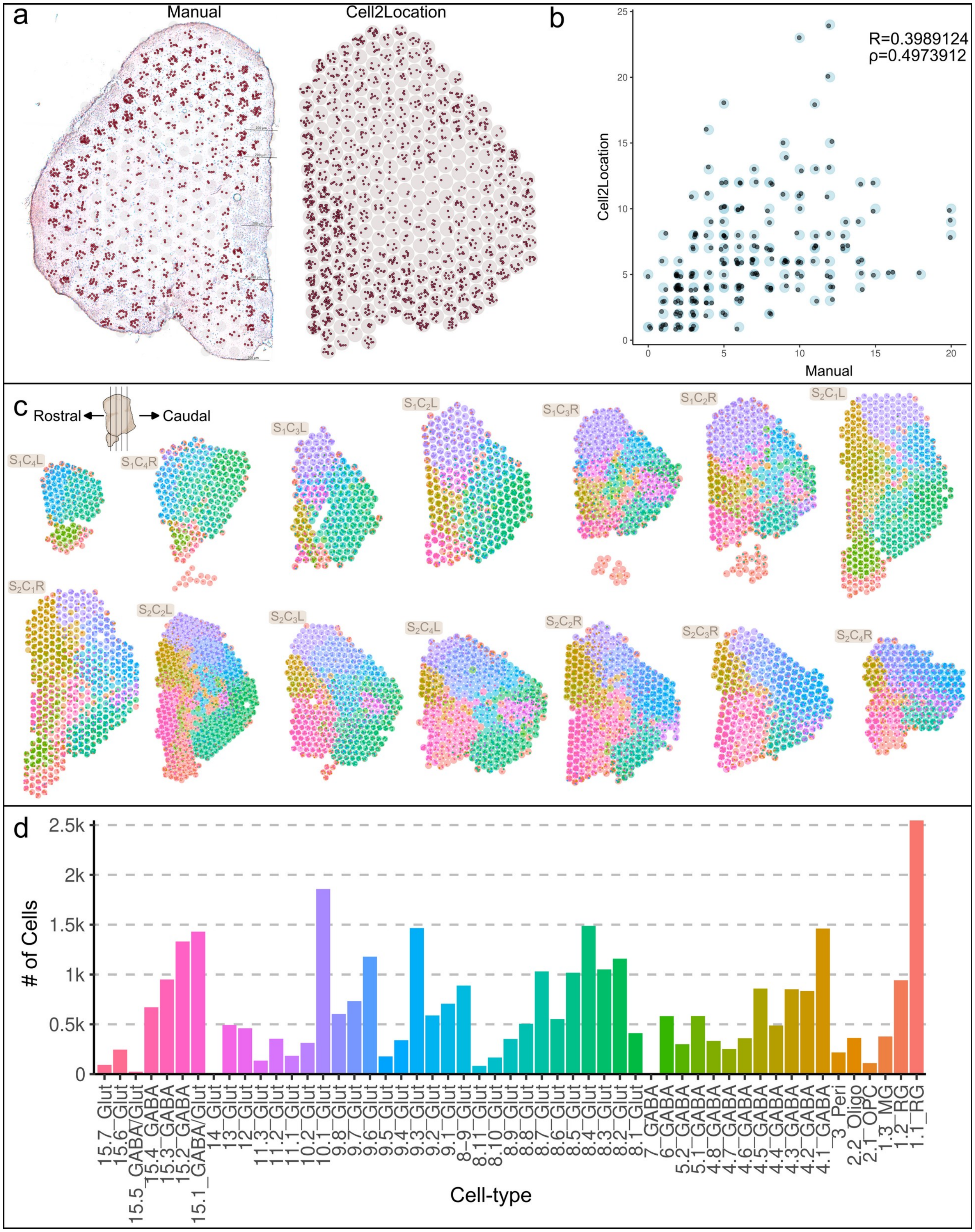


**Figure S7. Cell-type location and abundance prediction.** A. Comparison of manually counted estimates (left) of the number of cells per spot compared with computational estimates (right) of abundance for select spots in a select tissue hemisphere ($S_1C_2R$). Large circles represent spots and smaller dots within them represent estimated cells. B. Correlation of manual and computation estimate of number of cells for spots shown in the previous panel (Pearson's r=0.3989124; ρ=0.4973912). Points are jittered slightly, within the radius of the blue circles, in order to better visualize the number of spots driving the correlation. C. Cell2location cell-type abundance estimates in spots in all tissue hemispheres. D. Number of cells from each cell-type in all spots.

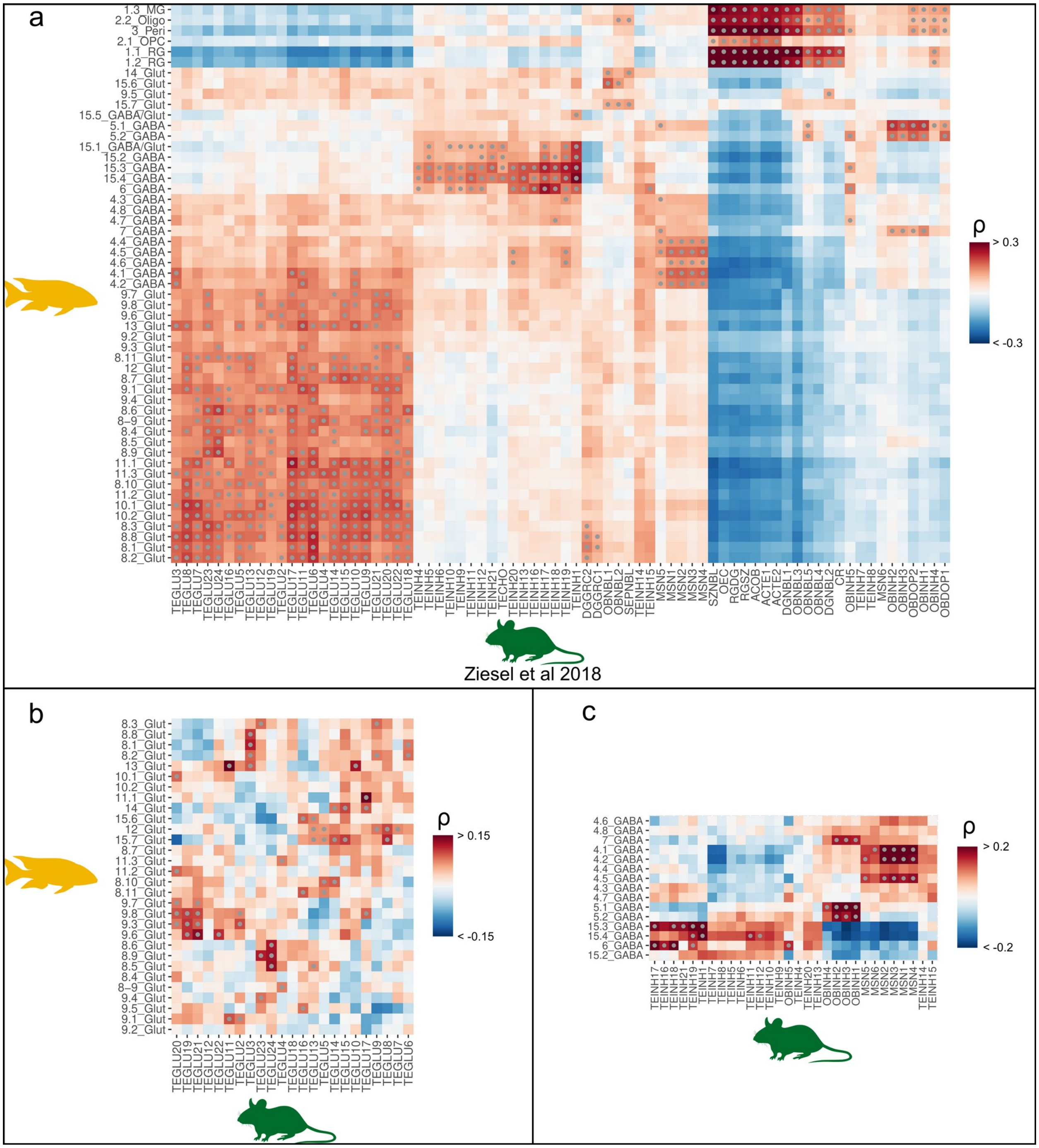


**Figure S8. Correlation gene expression profiles of shared marker genes between cichlids and mice.** A.Correlation of transcriptomic profiles by cell-type of DEGs shared between cichlids and mice. Gray dots denote cell-type pairs with correlations significant after permutation testing ($n_{perm}$=10,000). B-C. Correlations between glutamatergic and GABAergic cichlid and mouse cell-types, respectively.

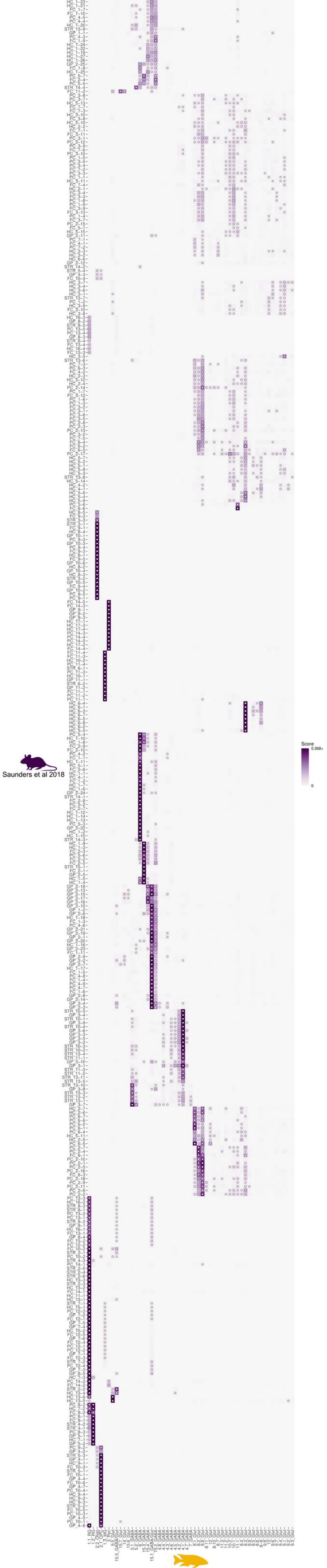
Saunders et al 2018
Score
0.368+
0

**Figure S9. Comparison of cichlid and mouse cell-types using SAMap.** Similarity score of cell-types in the cichlid and mouse telencephalon with dots denoting similarity scores greater than all permutations.

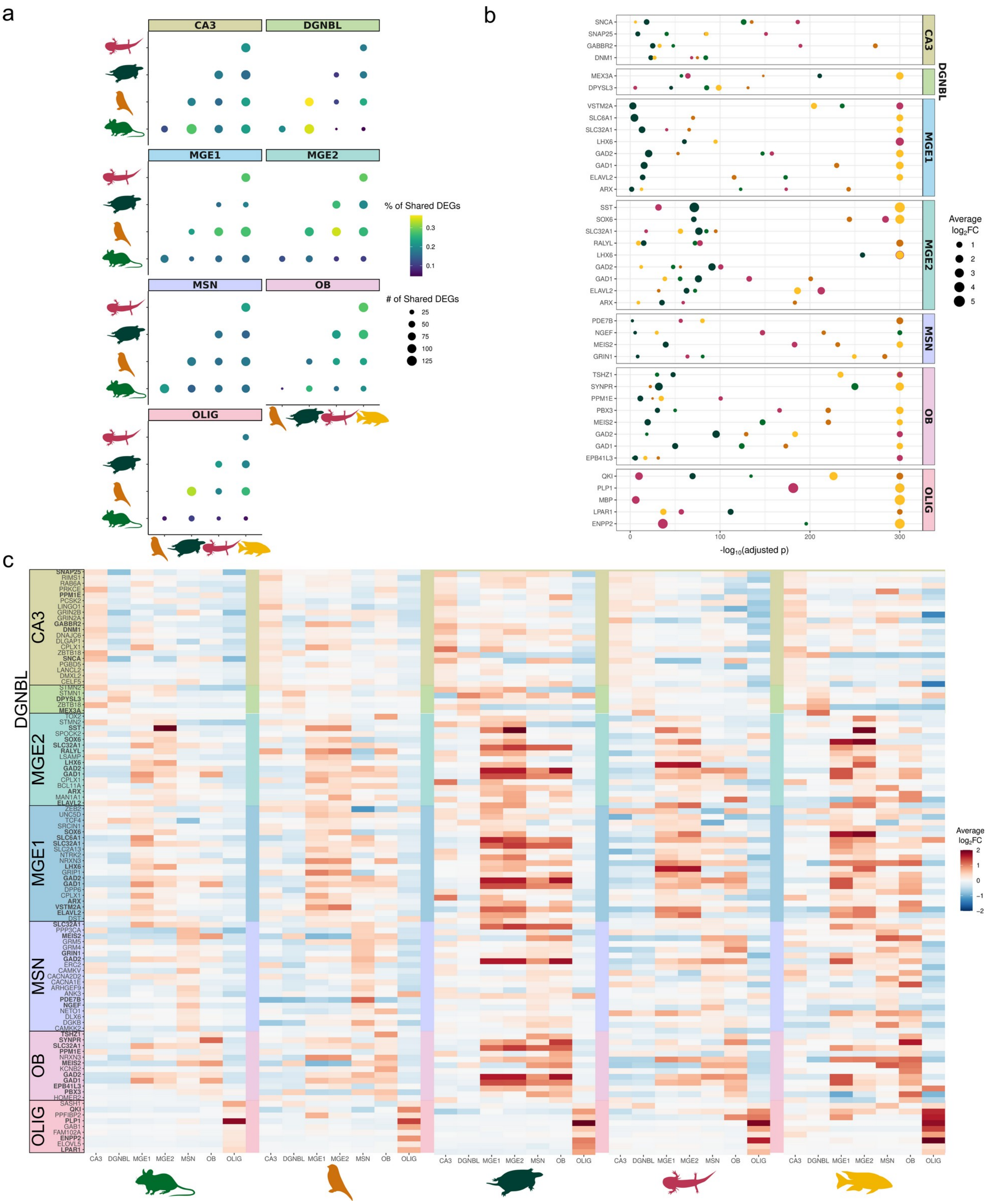


**Figure S10. Genes shared across conserved cell-types in vertebrates.** A. Number and percent of shared marker genes across pairwise combinations of vertebrates in strongly conserved cell-types in all vertebrates. Dot color signifies the percent of markers shared and dot size reflects the number of marker genes shared. B. Marker genes shared in all vertebrates in strongly conserved cell-types. Dot size signifies the average $log_2FC$ of the marker and the -$log_{10}$

of the adjusted p-value is shown on the x-axis. Dot color reflects the species of origin (green=mouse, brown=bird, teal=turtle, pink=axolotl and yellow=cichlid). C. Heatmap of average $log_2FC$ of marker genes (in bold text) and genes with positive log2FC in all vertebrates. Heatmaps are shown for each species separately using the same orthologous genes.

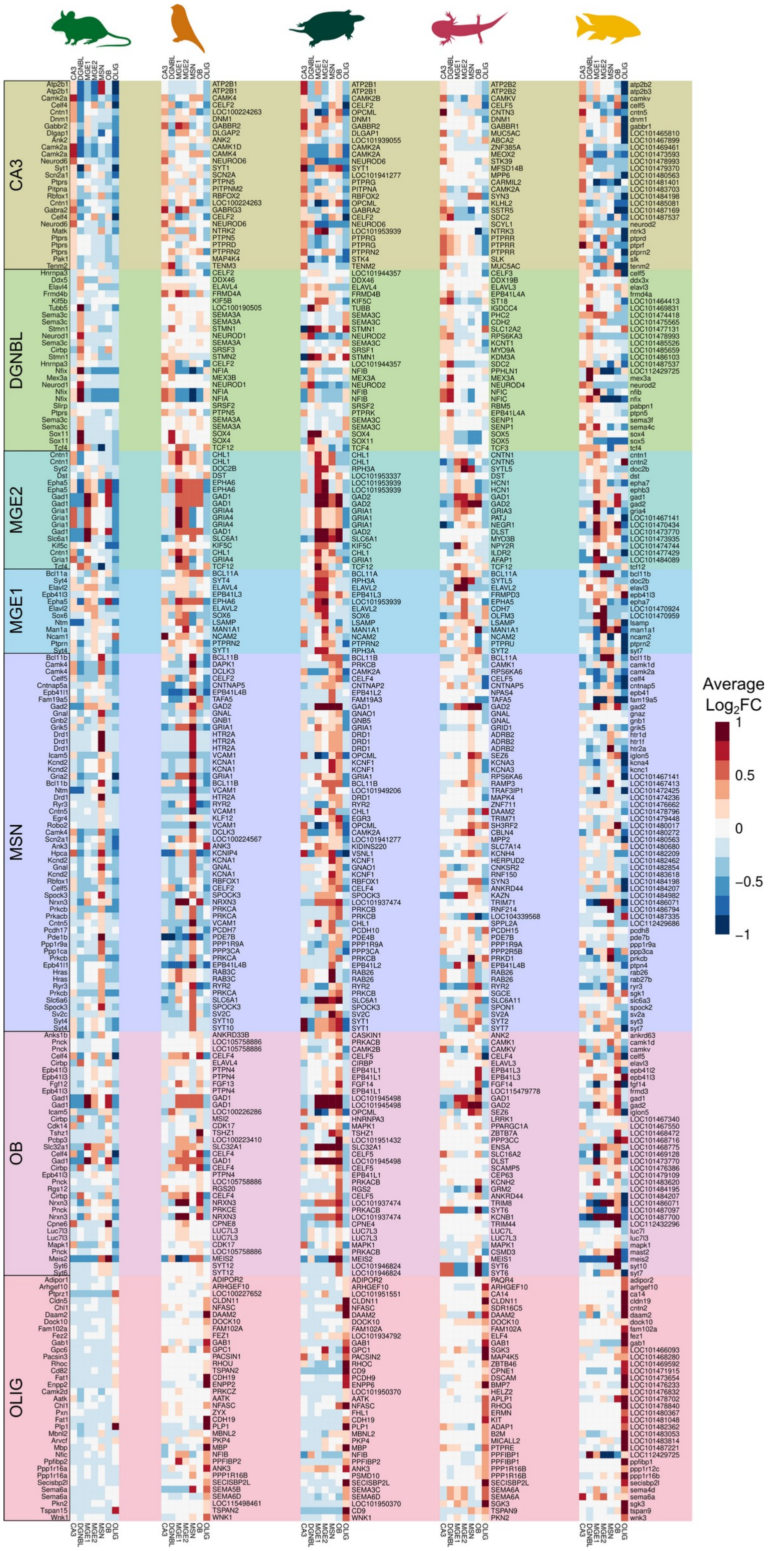


**Figure S11. Genes driving conserved cell-type relationships across vertebrates.** Heatmap of average $log_2FC$ of genes driving conserved cell-types in vertebrates. Heatmaps are shown for each species separately using genes orthologous to the same cichlid gene.

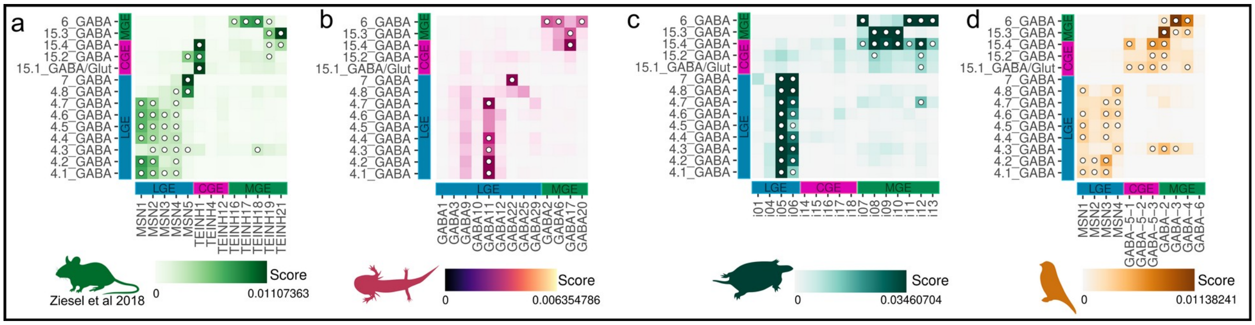


**Figure S12. Comparison of GABAergic cell-types across vertebrates.** A-D. Similarity scores of GABAergic cell-types in SAMap comparisons of cichlids to mice, axolotls, turtles, and birds respectively. Dots denote similarity scores greater than all permutations. Colored labels on the x-axis indicate the previously described origin of the cell-type (LGE, CGE and MGE). Colored labels on the y-axis indicate the putative origin of the cichlid cell-type.

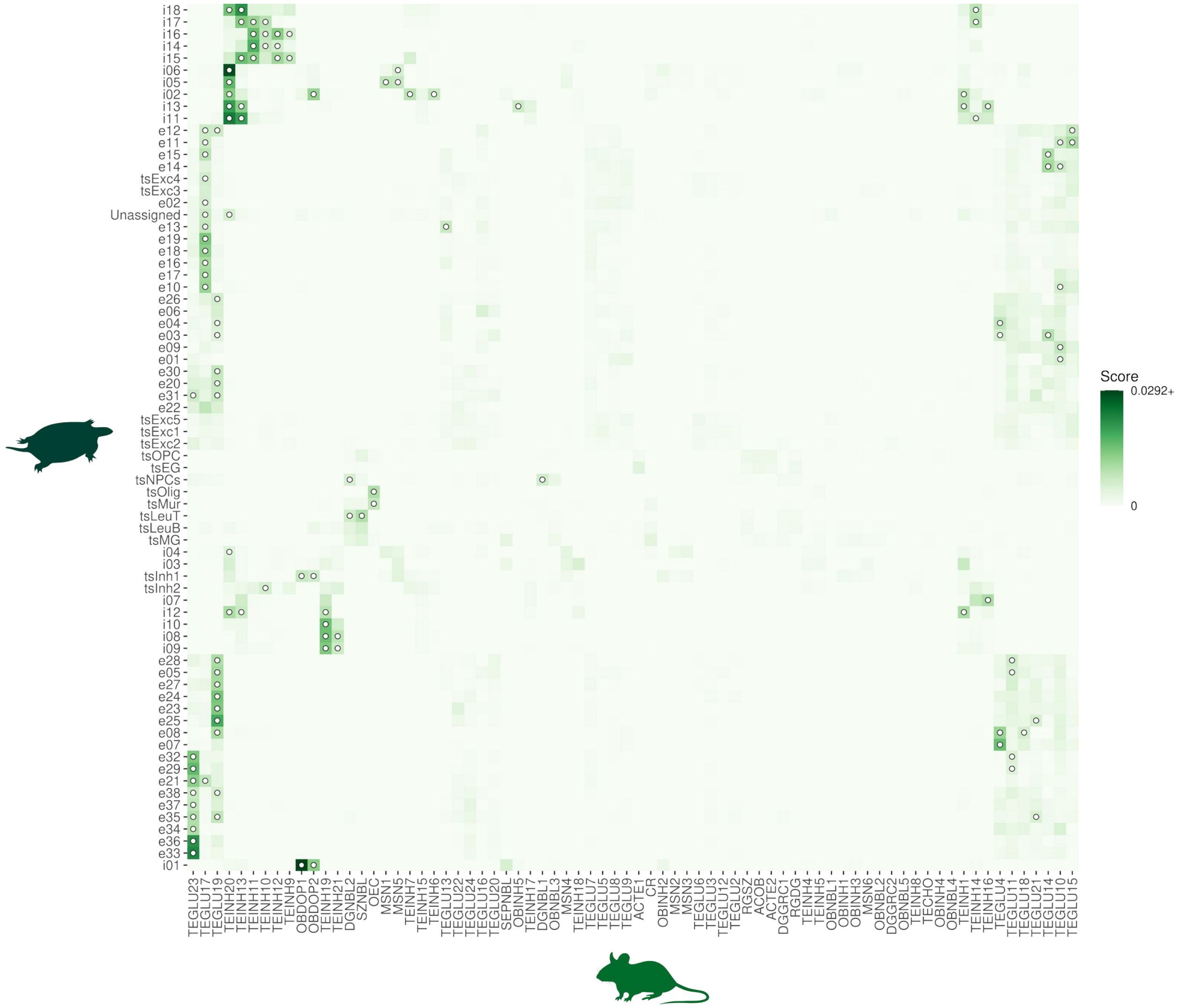


**Figure S13. Comparison of turtle and mouse cell-types using SAMap.** Similarity score of cell-types in the turtle and mouse telencephalon with dots denoting similarity scores greater than all permutations.

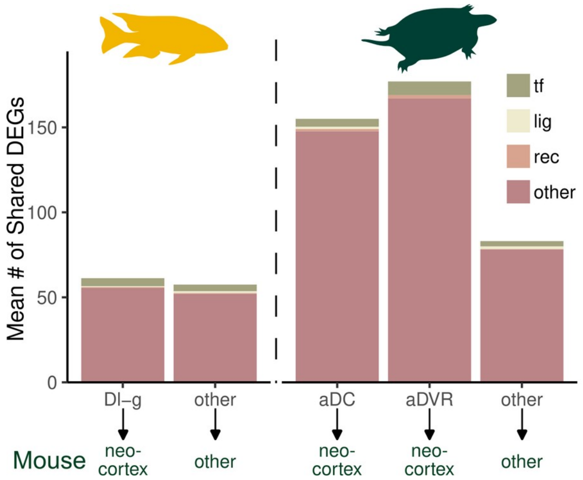


**Figure S14. Composition of conserved marker genes between cichlids and turtles with the mammalian neocortex.** The composition of conserved marker genes between significant cichlid-turtle and turtle-mouse cell-types were found by transcription factors, neuromodulatory ligands and receptors. The composition was compared between cichlid cell-types with significant transcriptional similarity to the neocortex (12_Glut with TEGLU6, and 12_Glut with TEGLU9) and other significant pairs. Likewise, for turtle aDC () and aDVR (e01 with TEGLU10, e07 with TEGLU4, e08 with TEGLU4, and e14 with TEGLU10) cell-types with the neocortex and other significant cell-type pairs.

**RESULTS**

Manual annotation of anatomical regions

Due to the slight angle of the cryosectioning plane, left and the right telencephalic hemispheres captured on each 10µm section differed from each other rostrocaudally. For this reason, each hemisphere was visualized and analyzed separately downstream irrespective of its neighboring hemisphere. Individual hemispheres of coronal sections were organized along the rostrocaudal axis based on tissue cytoarchitectural landmarks using atlases of the *Astotilapia burtoni* cichlid brain as references[1–3]. For each coronal hemisphere, anatomical region identities of spots were manually annotated using Loupe Browser visualization software (10X Genomics). 26 total anatomical regions were annotated based on 1) tissue cytoarchitecture via visual inspection of H&E stained brightfield images of the tissue sections (Fig. S1) and 2) previously established expression of well-studied genes in the subregions of the teleost telencephalon. Annotations of pallial and subpallial subdivisions differ considerably throughout teleost literature, therefore many decisions regarding the boundaries of these regions were based on shared transcriptional patterns revealed by unbiased clustering coupled with cytoarchitectural characteristics observed in H&E images of tissue sections. Within pallial and subpallial areas, further distinction between individual subregions was based on combinatorial gene expression patterns described below and shown in Fig. S6.

In the teleost telencephalon, dorsal/pallial regions and the olfactory bulb glomerular/mitral (or external) layers (OB gml) generally express glutamatergic markers (*slc17a6/vglut.2*, *slc17a7/vglut.1*(LOC101484681)) while ventral/subpallial regions and the olfactory bulb granule (or internal) cell layer (OB gc) are largely GABAergic (*gad1*, *gad2*)[3]. This trend is reflected in our spatial data. Ventral/subpallial areas (Vv, Vd, Vs, Vc, Vl) and the OB gcl expressed *dlx5* and *dlx2* (to a lesser extent)[4]. In the dorsal/pallial regions, we observed expected expression patterns of genes shown to be expressed in the teleost pallium, including *eomesa* (LOC101480282)*, neurod1, neurod6* (LOC101472195)*, emx1* (LOC101483575)*, and bhlhe22,* described below. We note that pallial markers *tbr1* and *emx3* were largely absent from spatial data in contrast with their abundant expression in MC snRNA-seq pallial cell-types. Of the pallial markers, *bhlhe22* was expressed across all dorsal/pallial regions (Dm, Dl, Dc, Dd, Dp)[5]. Spots situated along the periphery of the tissue which expressed strong markers of radial glia cells (RGC) were annotated as belonging to the ventricular zone (VZ), a layer of cells lining the ventricular surfaces (midline and outer brain margins) of the teleost telencephalon. We termed the vVZ as a subset of transcriptionally-distinct spots situated at the ventral ventricular surface.

In the *Astotilapia burtoni* cichlid pallium, several major telencephalic divisions, including the Dm, Dc, and Dl, are partitioned into distinct cell groups based on cytoarchitecture[1]. The Dc corresponds to several widely-spaced groups of large, migrated cells and the number of Dc subdivisions varies considerably in teleost species. In the *A. burtoni* telencephalon, five Dc subdivisions have been proposed[1], which we used as a guideline for distinguishing general Dc territories in the MC telencephalon, though ours differed slightly based on additional usage of transcriptional profiles to inform decisions. Within the Dm, we identified Dm-1 in the rostral-most coronal section ($S_1C_4$) and Dm-3 which persisted caudally at the dorsal edge of the pallium. Dm-2 (Dm-2r, Dm-2c) is proposed as a specialization in cichlids and positioned ventral to Dm-3[1].

Dm-2r corresponded strongly to ST cluster 15 and was identified based on its distinctive shape and ventromedial position below Dm-3/Dc-3 close to the midline, while Dm-2c was only identified in one hemisphere ($S_1C_2R$) as a tightly packed group of cells lateral to Dm-2r near Dd (Fig. S5). Expression domains of *cnr1* and *epha4* (LOC101472694) were similar to patterns observed in the zebrafish telencephalon with some exceptions[6]. *Cnr1* was strongly expressed in the Dm-1, Dm-2r, and Dd- a region which is not consistently annotated in teleost studies- while *epha4* demonstrated higher expression in Dl territories[6]. In line with observations in cichlids[1–3], we identified the Dd as a large, uniform group of cells neighboring the Dm and Dl at rostral levels and positioned dorsal to Dp at caudal levels. The Dl, which demonstrated strong expression of *ucn*[7], is a large, laterally positioned region with several distinct cell groups. The Dl-g, a group of tightly packed, small cells in the dorsal Dl, is another putative specialization in the cichlid brain, with other teleost Dl divisions generally being divided into a dorsal (dDl/Dl-d) and ventral (vDl/Dl-v) subdivisions[1]. Based on H&E images, the Dl-g was identified as largely corresponding to ST cluster 27 and was distinct from Dl-d and Dl-v due to its characteristic tightly packed cells and high expression of *slc17a7* (LOC101484681)[3,8,9], *pomc* (LOC101475601), and *pvalb7* (LOC101487165)[3,8,9,10]. In the area corresponding to the Dl-v, we observed a transcriptionally distinct population of tightly packed cells at its ventral-most pole, which we refer to as the Dl-vv in line with previous annotations in the *A. burtoni* telencephalon[2,11]. Some teleost studies identify a dense cell group in a similar area as the nucleus taenia (NT)[1]. The Dp displaced Dl-v at caudal telencephalic levels and demonstrated exclusive expression of *emx1*[3]. *Eomesa* demonstrated strongest expression in Dl-v/Dp regions with lower expression in areas corresponding to the Dl-g/Dd. *Neurod6* was expressed most strongly in Dm and Dl-d with lower expression in the Dd and Dp[5] while *neurod1* was expressed in the Dl-v and Dp[4].

Several divisions in the ventral telencephalon were generally easy to distinguish due to well-studied gene expression patterns and distinct cytoarchitectural features. The ventrally-located Vv was identified based on its expression of several genes, including *isl1* (LOC101475168)[4] and *nkx2-1*[4], and exclusive expression of *chat* (LOC101468574)[3], which marks cholinergic neurons. At caudal levels in the teleost telencephalon, a preoptic region is located in the ventral-most position along the midline, but due to the angling of the sectioning plane we believe that this region was not included in our sections. Dorsal to Vv, the Vd was divided into a rostral part (Vd-r) and caudal part (Vd-c) in line with observations in the *A. burtoni* telencephalon[1]. The Vs is positioned dorsal to the Vd and ventral to Dm along the midline and divided into lateral and medial parts, while a division referred to as the Vp appears below the Vs at caudal levels[1]. Instead, we chose to annotate the MC Vs as one unit and did not distinguish a separate Vp. We examined expression of several genes that mark principal striatal neurons (medium spiny neurons, MSNs) in the mammalian brain, including *penk* (LOC101474395), *tac1*, *six3*, and *sp9*[12], that have been studied in teleosts. Vd and Vs demonstrated high expression of *six3*[5], and the border between these regions was partially determined based on the distribution of *penk,* expressed strongly in the Vd*,* and *tac1*, which is more abundant in the Vs[6,13]. *Sp9* was expressed in subpallial territories with strongest expression in the Vd and Vc[5], which is a small, migrated group of cells lateral to Vv and Vd. This nucleus was easily identified in the H&E images of many sections and corresponded largely with ST cluster 6 (Fig. S5). In the cichlid subpallium, two other migrated groups of cells have been identified- the Vl, and Vi[1]. We observed strong co-location of *sst* (LOC101485677) and *npy* in the Vl[1415], the most lateral division of the ventral telencephalon, which additionally exhibited high expression of crh[7]. We only identified Vi in one caudal hemisphere ($S_2C_3R$) as a small cluster of cells located centrally in a largely cell-poor region of the subpallium that we refer to as the *Vx*. Additionally, we compared expression of dopaminergic genes (*th*, *slc6a3*, *slc18a2*) in ST data with patterns described in teleosts to further classify the Vd, Vc, and OB gc[1617] and observed expected expression of dopamine receptor *drd2* in the Vd, Vc, Vv, Vi, OB, and Dp[17].

Further determination of regional identities was based on expression patterns of transcriptional regulators. We looked at distribution of Zic family genes and observed *zic1* with strong expression in the Vv, *zic3* expressed in the Dl-v, Dp, Vv and Vc and *zic2* expressed in the Vv, Dp, and Dl-v[5]. LIM homeobox genes *lhx6* and *lhx8 (lhx7)* were expressed strongly in the Vv[18], while *lhx9* was localized in the OB gml[19] and Dp. Inhibitor of DNA Binding (ID) family members demonstrated expression patterns in line with observations in the adult teleost telencephalon[20]. *Id1*, which is expressed in radial glia cells (RGCs), was localized along the ventricular zone (VZ)[20]. Dm, Dc, and Dp regions demonstrated strong expression of *id2* while *id4* (LOC101483990) was expressed most strongly in the Vd and Vv, with lower expression in Dm-3 and other subpallial regions [20]. Furthermore, we observed localized expression of *vax1* in the Vc and *prdm12* and *prdm16* in the Vv, genes which are exclusively expressed in these locations in the zebrafish telencephalon[5].

<u>Comparison of manual and computational cell abundance estimation in spatial transcriptomics spots</u>
To determine the rough accuracy of computational cell abundance estimation performed by cell2location[21], cells were manually counted in a select tissue hemisphere ($S_1C_2R$) and compared (Supp F7). Generally, both methods identify a large density of cells near the peripheral edges of the tissue and fewer cells in the middle. The Pearson and Spearman correlation coefficients were calculated to quantify this relationship. There was moderate-high agreement as the Pearson correlation was 0.3989124 and the Spearman correlation was 0.4973912 (Supp F7B).

<u>Correlations of expression of common marker genes in cichlid and mouse cell-types</u>
Several consistent cell-type relationships were observed using both the SAMap approach and correlation approach (Supp F8), including the following. *Six3*+ 4_GABA cichlid cell-types in the Vd and Vs were strongly correlated with mouse striatal MSNs (MSN1-4). Cichlid inhibitory populations located in the olfactory bulb were strongly correlated with their mammalian counterparts (OBINH1-4). Cichlid 6_GABA and 15.3_GABA were significantly correlated with MGE interneurons (TEINH17-19 and TEINH21).

Cichlid excitatory neuronal populations located in the Dl-g (8.1_Glut, 8.2_Glut and 8.8_Glut) had the greatest and significant similarity to mouse cells from pyramidal layer 6 of the cortex. Additionally, another Dl-g cell-type (12_Glut) was similar to several neocortical populations (TEGLU7-9). Most cichlid Dl-v cell-types (8.3_Glut, 8.5_Glut, 8.6_Glut and 8.9_Glut) have significant correlations with excitatory neurons in the hippocampus (TEGLU23-24). Cichlid 11.1_Glut located in the Dp was significantly correlated with piriform pyramidal cells (TEGLU17).

SAMap comparison of cichlid cell-types to an additional mouse telencephalon scRNA-seq dataset

Another scRNA-seq dataset from the mouse telencephalon was used to determine the robustness of observed similarities between cichlid and mouse cell-types (Supp F9). In this comparison, the mouse dataset consisted of regions annotated as Frontal Cortex (FC), Globus Pallidus (GP), Hippocampus (HC), Posterior Cortex (PC) and Striatum (STR).

Cichlid 1.1_RG expresses many markers of neurogenesis such as *Sox4* and demonstrates transcriptional similarities to mouse cell-types with a class label of "Neurogenesis" given by Saunders et al. Some examples of mammalian neurogenic cell-types similar to cichlid 1.1_RG include: HC_13-1 (SGZ), HC_13-2 (SGZ), HC_13-3 (SGZ), HC_13-4 (SGZ), HC_13-6 (SGZ), STR_2-1 (RMS), STR_2-2 (RMS), STR_2-3 (RMS), STR_2-4 (RMS), STR_2-5 (RMS), STR_2-6 (RMS) and mitotic GP_4-6. As a *fapb7*+ radial glia cell-type, 1.1_RG shows transcriptional similarity to the *Fabp7*+ astrocyte cell-type HC_7-3. Cichlid 1.1_RG and cichlid 1.2_RG show transcriptional similarity to the following astrocyte cell-types: FC_8-1, FC_8-2, GP_5-1, GP_5-2, HC_7-1, HC_7-2, PC_8-1, PC_8-2 and PC_8-3. However 1.2_RG generally showed greater similarity to these astrocyte cell-types except for *Fabp7*+ FC_8-2.

Cichlid *tnr*+ 2.1_OPC has transcriptional similarity to *Tnr*+ polydendrocyte cell-types: PC_10-1, PC_10-3, PC_10-4, GP_4-1, GP_4-4, GP_4-5, GP_4-7, FC_10-1, FC_10-3, FC_10-5, FC_10-6, HC_9-1, HC_9-2, HC_9-3, HC_9-4, STR_5-1, STR_5-2 and STR_5-3. Cichlid 2.2_Oligo has transcriptional similarity to oligodendrocyte cell-types: PC_9-2, PC_9-3, PC_9-4, PC_9-5, FC_9-1, FC_9-3, FC_9-4, FC_9-5, GP_10-1, GP_10-2, GP_10-3, GP_10-4, GP_10-5, GP_10-6, HC_8-1, HC_8-2, HC_8-3, HC_8-4, STR_3-1, STR_3-2 and STR_3-3. Cichlid 1.3_MG has transcriptional similarity to microglia/macrophages: PC_11-1, PC_11-2, PC_11-3, PC_11-4, FC_11-1, FC_11-3, FC_11-4, GP_11-1, GP_11-2, HC_10-1, HC_10-2, STR_6-1 and STR_6-2. Cichlid 3_Peri has transcriptional similarity to cells from the endothelial tip: PC_14-3, PC_14-4, PC_14-5, FC_14-3, FC_14-4, FC_14-5, GP_9-1, GP_9-2, GP_9-3, HC_17-1, HC_17-2, HC_17-3 and HC_17-4.

Cichlid *sox4*+ 9.5_Glut has transcriptional similarity to neurogenic *Sox4*+ neurons from the hippocampus: HC_13-5 (SGZ) and HC_13-6 (SGZ). Cichlid 15.7_Glut was the reciprocal top hit of Cajal-Retzius cells in the frontal cortex.

This mouse dataset contains few Th+ dopaminergic inhibitory interneurons (STR_13-5 and STR_14-4), but cichlid 5.1_GABA and/or 5.2_GABA does have significant similarity to them, as well as to other striatal inhibitory interneurons. 6_GABA has transcriptional similarity to many *Pvalb*+ inhibitory interneurons: PC_5-1, PC_5-2, PC_5-3, FC_2-6, FC_2-7, FC_2-9, FC_2-10, HC_1-6 and HC_1-7. Additionally, 15.3_GABA shows transcriptional similarity to MGE-derived cell-types that were Npy+ (STR_15-1) and *Sst*+ (PC_5-6, FC_2-1, FC_2-2, FC_2-3, FC_2-4,

HC_1-3, HC_1-4, HC_1-5, HC_1-9, STR_15-1). 15.1_GABA/Glut, 15.2_GABA, and 15.4_GABA has transcriptional similarities to an Lhx6-enriched cell-type in the globus pallidus externus, as well as to many other globus pallidus cell-types. Cichlid inhibitory 4.1_GABA has significant transcriptional similarity to all of the following striatal spiny neurons, and several other cichlid has significant similarities to some: GP_3-1, GP_3-2, GP_3-3, GP_3-4, GP_3-5, GP_3-6, STR_10-1, STR_10-2, STR_10-3, STR_10-4, STR_10-5, STR_11-1, STR_11-2, and STR_11-3.

Cichlid excitatory neuronal populations located in the Dl-g (8.1_Glut, 8.2_Glut, 8.8_Glut, and 12_Glut) have the greatest and significant similarity to retrosplenial areas (PC_6-1, PC_6-2, PC_6-5, and PC_6-6), entorhinal cortex (PC_6-7 and HC_5-11) and other neocortical cell-types (PC_2-1, PC_2-2, PC_2-4, PC_2-5, PC_2-6, PC_2-7, PC_2-10, PC_2-11, PC_2-16 and PC_2-18). Cichlid 11.1_Glut located in the Dp has the greatest similarity to deep layer pyramidal cells (FC_6-6). Most cichlid Dl-v cell-types (8.3_Glut, 8.5_Glut and 8.6_Glut) have significant similarity to lateral CA3 principal cells (HC_6-3), but 8.3_Glut is the strongest. Additionally, 8.3_Glut shows strong similarity to many other hippocampal populations (HC_5-4, HC_5-5, HC_5-6, HC_5-9, HC_6-1, HC_6-2, HC_6-3, HC_6-4, HC_6-5, HC_6-6, HC_6-7 and HC_6-8).

<u>Genes driving effects between anatomical regions in the cichlid and mouse telencephalon</u>
Cichlid OB regions (OB gc, OB gml) demonstrated strong transcriptional similarities with the mouse main olfactory bulb (MOB) and genes that contributed to this pairing included *gpsm1*, identified as a novel marker of olfactory bulb granule cells in the mouse[22], and *arx*, which is critical for OB development and differentiation of *Th*+ neurons in this region in the mouse brain[23]. Other genes included markers of OB dopaminergic cells (*th, sall3*)[24] and genes involved in OB development and function (*dlx2, meis2, sp8, zic1, zic3*)[25].

Multiple cichlid regions mapped to the mouse nucleus accumbens (ACB) of the ventral striatum, including the Vd-c, Vd-r, and Vs. The vast majority of cells in the mammalian nucleus accumbens are medium spiny neurons (MSNs), and many genes that contributed to cichlid region-mouse ACB pairings are expressed in MSNs, including *bcl11b*, *baiap3*, *cpne5*, *id4*, *slc2a13*, *slc32a1*, *flrt3*, *hs6st2*, and *pbx3*[26]. Furthermore, genes enriched in specific MSN subtypes (D1 MSN-enriched *tac1* and D2 MSN-enriched *sp9*)[26] were also included in ACB pairings.

The cichlid Vv demonstrated strongest transcriptional similarity to the mouse lateral septal complex (LSX), as did the Vc. Several genes driving the Vv-LSX pairing are expressed in the adult murine lateral septum, including *nts*[27], *pnoc*[28], *nrxn3*[29] and *zic5*[30], while others have been shown to play important roles in the development of septal populations, including *isl1*[31], *prdm12*[32], *zic1*[33,34] and *zic3*[34]. The Vv additionally mapped to the mouse pallidum (PAL) along with the Vl. According to the Allen Mouse Brain Reference Atlas[35], the PAL contains regions such as the globus pallidus, medial septal complex, and the bed nuclei of the stria terminalis, among others. Several genes that contributed to the Vv-PAL pairing included *pvalb* and *zic1*, which are expressed in the adult mouse medial septum[33], and *lhx8*, which plays a role in development of medial septum cholinergic neurons which project to the hippocampus[36].

The mouse hippocampal CA3 region was transcriptionally similar to Dl-vv and two pallial regions in the cichlid rostral-most sections, the Dm-1 ($S_1C_4$) and Dl-d ($S_1C_4L$). Many genes driving the Dl-vv to CA3 pairing have been identified as enriched in the mouse hippocampal CA3 based on DNA microarray data (*grm1*, *bdnf*)[37] and single-cell RNA-seq (*grm1*, *bdnf*, *rasgrp1*, *ptk2b*, *prkce*, *pclo*, *cpe*, *spock1*, and *prkca*)[38].

The cichlid Dl-g mapped to the visual areas of the mouse isocortex (VIS). A subset of genes driving this comparison are reported as highly expressed in the mouse cerebral cortex (*camk2g*, *dusp6*, *egr1*, *gabra1*, *gadd45b*, *kcnf1*, *ncald*, *nrn1*)[39]. Furthermore, several genes driving the Dl-g to VIS pairing demonstrate layer-specific enriched expression in a transcriptomic atlas of mouse cortical layers (L2/3: *adgrb1*, *atp1a1*, *fosl2*, *nrn1*; L2/3 and L4: *golga7b*, *nr2f1*, *r3hdm1*; L5: *bhlhe40*, *camk2g*, *dusp6*, *gabra1*, *gfra2*; L6: *ncald*)[40].

### REFERENCES


1. Burmeister, S. S., Munshi, R. G. & Fernald, R. D. Cytoarchitecture of a cichlid fish telencephalon. *Brain Behav. Evol.* **74**, 110–120 (2009).
2. Munchrath, L. A. & Hofmann, H. A. Distribution of sex steroid hormone receptors in the brain of an African cichlid fish, Astatotilapia burtoni. *J. Comp. Neurol.* **518**, 3302–3326 (2010).
3. Maruska, K. P., Butler, J. M., Field, K. E. & Porter, D. T. Localization of glutamatergic, GABAergic, and cholinergic neurons in the brain of the African cichlid fish, Astatotilapia burtoni. *J. Comp. Neurol.* **525**, 610–638 (2017).
4. Ganz, J. *et al.* Subdivisions of the adult zebrafish subpallium by molecular marker analysis. *J. Comp. Neurol.* **520**, 633–655 (2012).
5. Diotel, N. *et al.* Comprehensive expression map of transcription regulators in the adult zebrafish telencephalon reveals distinct neurogenic niches. *J. Comp. Neurol.* **523**, 1202–1221 (2015).
6. Aoki, T. *et al.* Imaging of neural ensemble for the retrieval of a learned behavioral program. *Neuron* **78**, 881–894 (2013).
7. Website. Grone, B. P., Butler, J. M., Wayne, C. R. & Maruska, K. P. Expression patterns and evolution of urocortin and corticotropin-releasing hormone genes in a cichlid fish. Journal of Comparative Neurology vol. 529 2596–2619 Preprint at https://doi.org/10.1002/cne.25113.
8. Hu, C. K. *et al.* Identification of prohormones and pituitary neuropeptides in the African cichlid, Astatotilapia burtoni. *BMC Genomics* **17**, 660 (2016).
9. Mueller, T., Dong, Z., Berberoglu, M. A. & Guo, S. The dorsal pallium in zebrafish, Danio rerio (Cyprinidae, Teleostei). *Brain Res.* **1381**, 95–105 (2011).
10. Furlan, G. *et al.* Life-Long Neurogenic Activity of Individual Neural Stem Cells and Continuous Growth Establish an Outside-In Architecture in the Teleost Pallium. *Curr. Biol.*

**27**, 3288–3301.e3 (2017).

11. Huffman, L. S. *et al.* Distribution of nonapeptide systems in the forebrain of an African cichlid fish, Astatotilapia burtoni. *J. Chem. Neuroanat.* **44**, 86–97 (2012).
12. Xu, Z. *et al.* SP8 and SP9 coordinately promote D2-type medium spiny neuron production by activating expression. *Development* **145**, (2018).
13. Ogawa, S. *et al.* Cloning and expression of tachykinins and their association with kisspeptins in the brains of zebrafish. *J. Comp. Neurol.* **520**, 2991–3012 (2012).
14. Porter, D. T., Roberts, D. A. & Maruska, K. P. Distribution and female reproductive state differences in orexigenic and anorexigenic neurons in the brain of the mouth brooding African cichlid fish, Astatotilapia burtoni. *J. Comp. Neurol.* **525**, 3126–3157 (2017).
15. Pickavance, L. C., Staines, W. A. & Fryer, J. N. Distributions and colocalization of neuropeptide Y and somatostatin in the goldfish brain. *J. Chem. Neuroanat.* **5**, 221–233 (1992).
16. Yamamoto, K., Ruuskanen, J. O., Wullimann, M. F. & Vernier, P. Differential expression of dopaminergic cell markers in the adult zebrafish forebrain. *J. Comp. Neurol.* **519**, 576–598 (2011).
17. O’Connell, L. A., Fontenot, M. R. & Hofmann, H. A. Characterization of the dopaminergic system in the brain of an African cichlid fish, Astatotilapia burtoni. *J. Comp. Neurol.* **519**, 75–92 (2011).
18. Mueller, T., Wullimann, M. F. & Guo, S. Early teleostean basal ganglia development visualized by zebrafish Dlx2a, Lhx6, Lhx7, Tbr2 (eomesa), and GAD67 gene expression. *J. Comp. Neurol.* **507**, 1245–1257 (2008).
19. Alunni, A. *et al.* Cloning and developmental expression patterns of Dlx2, Lhx7 and Lhx9 in the medaka fish (Oryzias latipes). *Mech. Dev.* **121**, 977–983 (2004).
20. Diotel, N., Beil, T., Strähle, U. & Rastegar, S. Differential expression of id genes and their potential regulator znf238 in zebrafish adult neural progenitor cells and neurons suggests

distinct functions in adult neurogenesis. *Gene Expr. Patterns* **19**, 1–13 (2015).

21. Kleshchevnikov, V. *et al.* Cell2location maps fine-grained cell types in spatial transcriptomics. *Nat. Biotechnol.* **40**, 661–671 (2022).
22. Edsgärd, D., Johnsson, P. & Sandberg, R. Identification of spatial expression trends in single-cell gene expression data. *Nat. Methods* **15**, 339–342 (2018).
23. Yoshihara, S.-I., Omichi, K., Yanazawa, M., Kitamura, K. & Yoshihara, Y. Arx homeobox gene is essential for development of mouse olfactory system. *Development* **132**, 751–762 (2005).
24. Heng, X. *et al.* Sall3 correlates with the expression of TH in mouse olfactory bulb. *J. Mol. Neurosci.* **46**, 293–302 (2012).
25. Pignatelli, A. & Belluzzi, O. Neurogenesis in the Adult Olfactory Bulb. in *The Neurobiology of Olfaction* (ed. Menini, A.) (CRC Press/Taylor & Francis).
26. Gokce, O. *et al.* Cellular Taxonomy of the Mouse Striatum as Revealed by Single-Cell RNA-Seq. *Cell Rep.* **16**, 1126–1137 (2016).
27. Risold, P. Y. & Swanson, L. W. Chemoarchitecture of the rat lateral septal nucleus. *Brain Res. Brain Res. Rev.* **24**, 91–113 (1997).
28. Ikeda, K. *et al.* Distribution of prepro-nociceptin/orphanin FQ mRNA and its receptor mRNA in developing and adult mouse central nervous systems. *J. Comp. Neurol.* **399**, 139–151 (1998).
29. Uchigashima, M., Cheung, A., Suh, J., Watanabe, M. & Futai, K. Differential expression of neurexin genes in the mouse brain. *J. Comp. Neurol.* **527**, 1940–1965 (2019).
30. Mühlfriedel, S., Kirsch, F., Gruss, P., Chowdhury, K. & Stoykova, A. Novel genes differentially expressed in cortical regions during late neurogenesis. *Eur. J. Neurosci.* **26**, 33–50 (2007).
31. Developmental specification of forebrain cholinergic neurons. *Dev. Biol.* **421**, 1–7 (2017).
32. Rajendram, R., Preedy, V. R., Patel, V. B. & Martin, C. R. *The Neurobiology, Physiology,*

*and Psychology of Pain*. (Academic Press, 2021).

33. Wei, B. *et al.* The onion skin-like organization of the septum arises from multiple embryonic origins to form multiple adult neuronal fates. *Neuroscience* **222**, 110–123 (2012).
34. Inoue, T., Ota, M., Ogawa, M., Mikoshiba, K. & Aruga, J. Zic1 and Zic3 regulate medial forebrain development through expansion of neuronal progenitors. *J. Neurosci.* **27**, 5461–5473 (2007).
35. Available from mouse.brain-map.org.Allen Institute for Brain Science (2011).
36. Zhao, Y. *et al.* The LIM-homeobox gene Lhx8 is required for the development of many cholinergic neurons in the mouse forebrain. *Proc. Natl. Acad. Sci. U. S. A.* **100**, 9005–9010 (2003).
37. Bonaventure, P. *et al.* Nuclei and subnuclei gene expression profiling in mammalian brain. *Brain Res.* **943**, 38–47 (2002).
38. Cembrowski, M. S., Wang, L., Sugino, K., Shields, B. C. & Spruston, N. Hipposeq: a comprehensive RNA-seq database of gene expression in hippocampal principal neurons. *Elife* **5**, e14997 (2016).
39. Lein, E. S. *et al.* Genome-wide atlas of gene expression in the adult mouse brain. *Nature* **445**, 168–176 (2006).
40. Belgard, T. G. *et al.* A transcriptomic atlas of mouse neocortical layers. *Neuron* **71**, 605–616 (2011).
41. Butler, A. B. & Hodos, W. *Comparative Vertebrate Neuroanatomy: Evolution and Adaptation*. (John Wiley & Sons, 2005).
