## Supplementary material for "Spatially resolved cell atlas of the teleost telencephalon and deep homology of the vertebrate forebrain": Inventory of Supplement

Supplementary Dataset 1.
Summary data for subjects (Sheet 1) and sequencing information for each capture area (Sheet 2).

Supplementary Dataset 2.
Silhouette scores of bootstraps of the data from ChooseR, including the 95% confidence interval and median of the silhouette scores. These scores are shown all combinations of parameters tested and the number of resulting clusters.

Supplementary Dataset 3.
Summary data of unbiased clusters, including the highly variable genes that formed the clusters (Sheet 1), the number of spots from each cluster in each tissue hemisphere (Sheet 2), the top 5 markers of clusters (Sheet 3) and all cluster markers (Sheet 4).

Supplementary Dataset 4.
Differentially expressed genes in manually annotated anatomical regions of the teleost telencephalon.

Supplementary Dataset 5.
Cell-type abundance estimates by cell2location in each spot.

Supplementary Dataset 6.
Results from SAMap comparative analysis of cell-types in the cichlid and mouse telencephalon. The similarity score and p-value from permutations of the data are provided.

Supplementary Dataset 7.
Conserved marker genes (Sheet 1) and genes driving effects (Sheet 2) in significant cell-pairs in the cichlid and mouse telencephalon.

Supplementary Dataset 8.
Results from SAMap comparative analysis of cell-types in the axolotl (Sheet 1), turtle (Sheet 2) and songbird (Sheet 3) forebrain with the cichlid telencephalon.

Supplementary Dataset 9.
Conserved marker genes in significant cell-pairs in the axolotl (Sheet 1), turtle (Sheet 2) and songbird (Sheet 3) forebrain with the cichlid telencephalon. Genes driving effects in significant cell-pairs in the axolotl (Sheet 4), turtle (Sheet 5) and songbird (Sheet 6) forebrain with the cichlid telencephalon.

Supplementary Dataset 10.

Conserved marker genes and genes with positive logFC values without requiring differential expression (Sheet 1) and genes driving effects (Sheet 2) in strongly conserved cell-types in all vertebrates.

Supplementary Dataset 11.
Results from SAMap comparative analysis of brain regions in the turtle forebrain (Sheet 1) and mouse telencephalon (Sheet 2) with the cichlid telencephalon.

Supplementary Dataset 12.
Conserved marker genes in significant brain region pairs in the turtle forebrain (Sheet 1) and mouse telencephalon (Sheet 2) with the cichlid telencephalon. Genes driving effects in significant brain region pairs in the turtle forebrain (Sheet 1) and mouse telencephalon (Sheet 2)

Supplementary Dataset 13.
Composition of genes driving significant cell-type pairs in neocortical-like structures by gene categories in the turtle, cichlid and mouse forebrain (Sheet 1). Composition of genes driving significant brain region pairs in neocortical-like structures by gene categories in the turtle, cichlid and mouse forebrain (Sheet 2). Gene categories for which the composition was examined (Sheet 3). These categories include: transcription factors, neuromodulatory ligands and receptors.
